# Dynamic Reprogramming of Fungal Cell Walls Underlies Germination and Immune Exposure in Zygomycetous Fungal Pathogens

**DOI:** 10.64898/2026.05.12.724644

**Authors:** Isha Gautam, Maaike Suuring, Ankur Ankur, Alicia Withrow, Yifan Xu, Jean-Paul Latgé, Benoit Briard, Tuo Wang

## Abstract

Fungal germination is a critical developmental transition that underlies environmental adaptation and pathogenicity, yet how the cell wall is molecularly reprogrammed during this process remains poorly understood. Here we show that germination of *Rhizopus delemar* involves a developmentally programmed transition from a β-1,3-glucan-rich dormant scaffold to a chitin-chitosan-dominated polarized wall. Using solid-state nuclear magnetic resonance spectroscopy and cytochemistry approaches, we show that resting conidia contains a rigid β-1,3-glucan- and chitosan-rich core beneath a persistent melanin layer. During swelling, this architecture is largely maintained, but germ tube emergence triggers complete shutdown of β-1,3-glucan synthesis and extensive chitin-chitosan enrichment. Distinct chitosan polymorphs are selectively enriched, while mobile polysaccharides are progressively incorporated into the rigid scaffold. This remodeling enhances neutrophil recognition of swollen and germinating conidia. Our study reveals a molecular mechanism linking fungal morphogenesis, cell wall remodeling, and morphotype-specific immune exposure during mucormycosis.

## INTRODUCTION

Fungal germination represents a critical developmental transition in which environmentally resilient, dormant spores are converted into metabolically active and invasive growth forms^1-4^. In pathogenic Mucorales, such as *Rhizopus delemar*, a major causative agent of mucormycosis, this process is tightly linked to virulence and disease progression, as the establishment of polarized growth enables rapid tissue invasion^5-7^. Dynamic remodeling of the cell wall during germination also modulates immune recognition^8-11^. Understanding how this architecture reorganizes throughout germination is therefore essential for deciphering the mechanisms underlying pathogenicity, antifungal resistance, and morphogenetic regulation^12,13^.

Unlike ascomycete fungi, mucoralean fungi possess distinctive biochemical and structural features, most notably a high abundance of chitosan, a partially deacetylated derivative of chitin that significantly influences wall mechanics, permeability, and immune recognition^14^. During germination, dormant conidia undergo isotropic swelling followed by polarized germ tube emergence, processes that require coordinated cell wall biosynthesis and remodeling of structural polysaccharides^15,16^. However, the molecular-level organization, polymorphism, and dynamic partitioning of these polymers between rigid structural scaffolds and mobile matrix components throughout germination remain poorly understood in mucoralean cells. Cell wall constituents serve not only as essential structural elements but also as key pathogen-associated molecular patterns (PAMPs); consequently, developmental remodeling of wall architecture may directly influence innate immune recognition and host responses. Neutrophils and macrophages are both recognized as important effectors against aerial fungal pathogens^17^. While it is established that *Rhizopus* species can subvert the physiological killing mechanisms of macrophages, the role of neutrophils during Rhizopus infection remains less understood^18^.

To address these knowledge gaps, we apply cellular solid-state NMR (ssNMR) spectroscopy and cell biology approaches to investigate cell wall remodeling in *R. delemar* across conidial germination and its interaction with human neutrophils^19-22^. Coupled with a novel suite of metabolic ^13^C labeling schemes that distinguish pre-existing from neo-synthesized material, this approach enables time-resolved investigation of biosynthetic remodeling during fungal development. This strategy enables us to define how major wall polysaccharides are redistributed during the transition from dormant conidia to polarized germlings and how these developmental changes will be perceived by the host innate immunity. This work provides molecular-level insight into mucoralean cell wall remodeling and establishes a framework linking germination dynamics to structural adaptation and host-pathogen interactions.

## RESULTS

### Resting conidia of *R. delemar* exhibits a β-glucan- and chitosan-rich cell wall architecture

The cell wall of the resting conidium exhibited three morphologically distinct zones (**Figure 1A, B**): (i) an electron-dense outer layer, (ii) a granular electron-dense middle layer and (iii) an electron-lucent inner layer adjacent to the plasma membrane (**Figure 1B**, left). The granular middle layer appeared electron-lucent when samples were fixed without osmium tetroxide (OsO_4_) and embedded in hydrophilic LR White resin to minimize protein and lipid oxidation (**Figure 1B**, right). Accordingly, the outermost electron-dense layer of the conidium is identified as melanin-rich.

**Figure 1.**
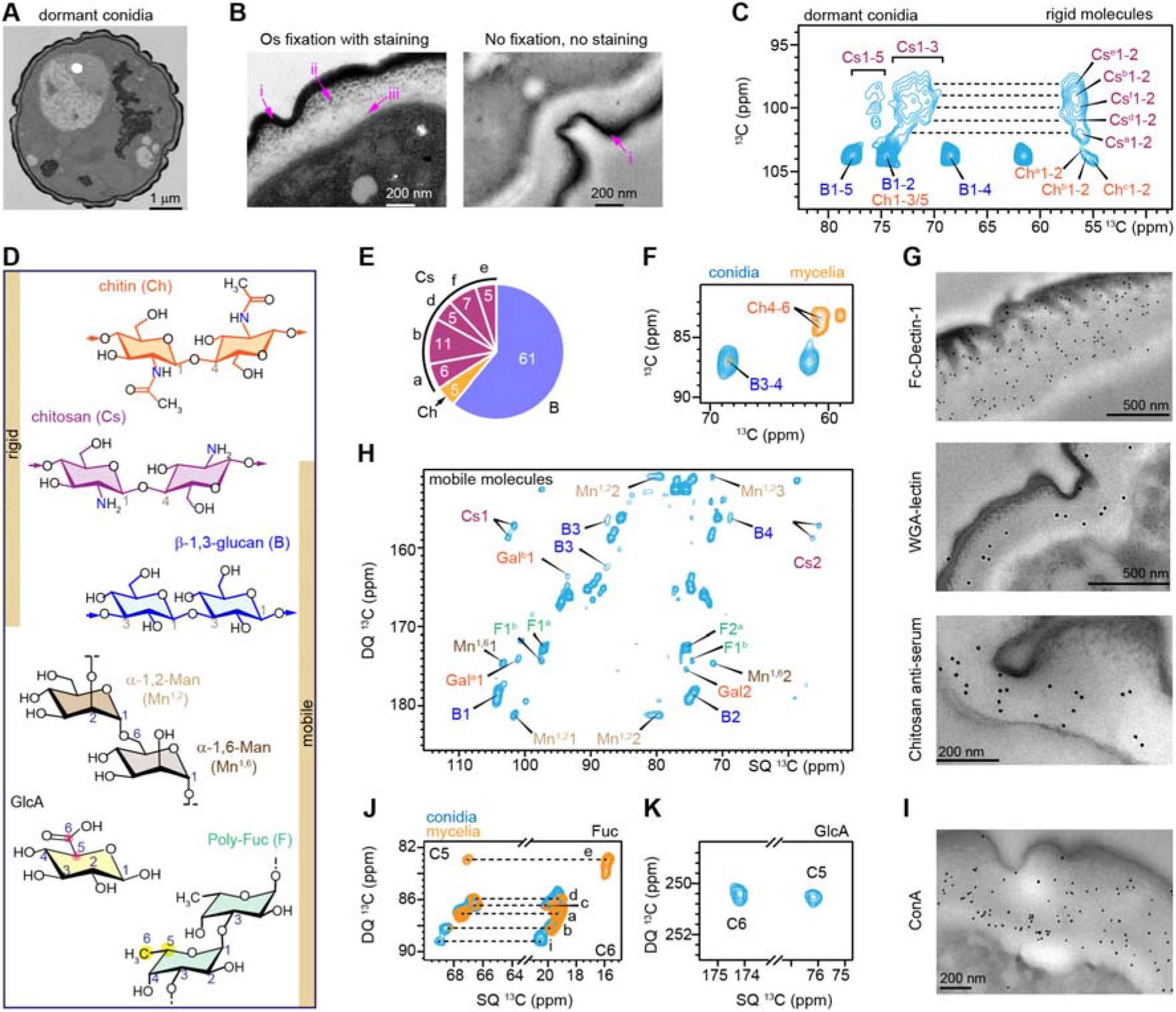
Dynamics and structure of polysaccharides in the *Rhizopus* conidial cell wall. (**A**) TEM images of the dormant conidia reveal a multilayered cell wall architecture, including an electron-dense outer layer and inner polysaccharide-rich regions. (**B**) At high magnification, osmium-fixed, epoxy-embedded samples (left) preserve structural integrity, whereas LR White–embedded samples prepared without OsO_4_ (right) enhance contrast in the outer wall. Sub-labels denote (i) outer electron-dense region, (ii) granular region, and (iii) electron-lucent inner layer. Without OsO_4_, the outer layer appears as a thin electron-lucent band consistent with melanin, with the remaining regions largely electron-lucent. (**C**) Rigid molecules in the dormant conidial cell walls were detected by 2D ^13^C-^13^C CORD spectra, showing signals from β-1,3-glucan (B), chitin (Ch), and chitosan (Cs). Superscripts indicate different structural forms of each polysaccharide, and the numbers denote carbon positions. For example, Cs^a^1-2 represents the correlation between carbons 1 and 2 in type-a chitosan. (**D**) Structural representations of carbohydrates. Yellow bars indicate whether each molecule is detected in the rigid phase, the mobile phase, or in both. NMR abbreviations are given. Key carbon sites are numbered. (**E**) Composition of rigid carbohydrates in dormant conidia based on intensity analysis of the 2D CORD spectrum. (**F**) Overlay of 2D CORD spectra of dormant conidia (blue) and 3-day-old mycelia (orange). (**G**) Cytochemical labeling of dormant conidia shows gold-specific staining for β-1,3-glucan with Fc-Dectin-1, chitin (with Wheat germ agglutinin lectin (WGA-lectin), and anti-chitosan rabbit antiserum. Black dots represent electron-dense gold particles indicating probe binding. Scale bars: 500 nm. (**H**) Mobile molecules in dormant conidia detected by the 2D ^13^C-DP refocused J-INADEQUATE spectrum. In addition to β-1,3-glucan and two forms of chitosan, signals are detected for Fuc (F), Gal, α-1,2-Man (Mn^1,2^), and α-1,6-Man (Mn^1,6^). (**I**) Cytochemical binding of Concanavalin-A lectin (ConA) to dormant conidial cell walls confirms the presence of mannans and/or glycoproteins. Zoomed-in view of 2D ^13^C-DP J-INADEQUATE spectrum was presented for (**J**) C5-C6 correlations of polymeric fucoses and (**K**) glucuronic acid (GlcA). Polymeric fucoses show five forms (a,b, c, d, i) as traced by dashed lines in dormant conidia (blue) while type-e is present only in mycelia (orange).

SsNMR spectra collected on intact dormant *R. delemar* conidia revealed signals from chitin, chitosan, and β-1,3-glucan within the rigid core, as selectively detected by 2D ^13^C-^13^C correlation experiments using dipolar-coupling-based ^1^H-^13^C cross polarization (**Figures 1C, D**; **Supplementary Fig. 1**). The crystalline domain formed by chitin and chitosan showed extensive structural polymorphism, resolving into five chitosan types (Cs^a^, Cs^b^, Cs^d-f^) and three chitin forms (Ch^a-c^; **Figure 1C**). Intensity analysis indicated that linear β-1,3-glucan constitutes 61% of the rigid molecules, followed by chitosan, whose five forms account for more than one-third of the dormant conidial wall (**Figure 1D, E**). In contrast, chitin contributes only 5% of the rigid phase (**Figure 1E; Supplementary Fig. 2**). This β-1,3-glucan-enriched conidial architecture contrasts sharply with the rigid core of *R. delemar* mycelial walls, which is predominantly composed of chitin and chitosan (**Figure 1F**)^14^. Cytochemical analysis further confirmed that β-1,3-glucan, chitin, and chitosan occupy a central inner position of the cell wall beneath the pigment layer, where they appear interwoven without clear spatial segregation (**Figure 1G**).

β-1,3-glucan and two distinct forms of chitosan were found to be distributed across both the rigid and mobile phases (**Figure 1H**), as evidenced by their signals in 2D ^13^C spectra acquired using direct ^13^C polarization with short recycle delays, which preferentially detect flexible species with rapid spin-lattice relaxation (**Figure 1D**; **Supplementary Fig. 3**). The mobile matrix also contains α-1,6- and α-1,2-linked mannose residues, characteristic of mannan backbones and branches (**Figure 1H**). Consistently, the inner cell wall layer was positively labeled with ConA-gold, indicating enrichment in mannan polymers and/or mannosylated proteins (**Figure 1I**; **Supplementary Fig. 3**).

Additionally, five structural forms of α-linked fucoses were resolved through their unique C5-C6 (methyl) signals (**Figure 1J**; **Supplementary Fig. 4**). Among these, type-b corresponds to α-1,3-linked fucose, likely forming the backbone of polymeric fucose, as previously observed in the mycelia of *R. delemar, R. oryzae* and *Phycomyces blakesleeanus*^14,23^. Glucuronic acid (GlcA), absent from our recent analyses of 3-day-old *R. delemar* mycelia, was clearly detected in dormant conidia through its characteristic C5-C6 (carbonyl) linkage (**Figure 1K**; **Supplementary Fig. 5**). This observation is consistent with previous reports across multiple mucoralean species and suggests morphotype-specific incorporation of GlcA-containing wall components^23,24^.

These spectroscopic and microscopic observations collectively demonstrate that the conidial cell wall of *R. delemar* exhibits a molecular architecture that is markedly distinct from those of the mycelial morphotype. The rigid phase is dominated by β-1,3-glucan and chitosan, with chitin present at low abundance, whereas β-1,3-glucan additionally extends into a mobile matrix enriched in mannan and polymers of fucose and glucuronic acid.

### Eumelanin compensates for unusually hydrophilic inner cell wall in dormant conidia

The low abundance of chitin, which forms water-excluding microfibrils, together with the high levels of β-1,3-glucan and chitosan, both of which readily interact with water, in dormant *R. delemar* conidia was unexpected, given the presumed need for high hydrophobicity to maintain environmental resilience during dormancy in dry environment. This unusual cell wall composition may be compensated by the outermost electron-dense eumelanin layer and its associated proteins.

To test this hypothesis, we subjected conidia to two complementary treatments. First, loosely associated molecules, proteins, and lipids were removed by boiling Sodium dodecyl sulfate (SDS) and dithiothreitol (DTT) extraction. This treatment preserved an NMR polysaccharide fingerprint nearly identical to that of intact conidia (**Supplementary Fig. 6**), confirming that the core wall architecture remained unchanged. Second, eumelanin formation was inhibited using bathocuproine disulfonate (BCS), which blocks tyrosinase/laccase activity required for eumelanin biosynthesis in zygomycetes^25-27^, resulting in the production of white conidia lacking melanin (**Supplementary Fig. 7-8**).

Analysis of the unpigmented conidia revealed a striking 3-4-fold increase in chitin abundance compared to intact conidia and the isolated cell wall fraction (**Figure 2A, B**). In contrast, the level of β-1,3-glucan in the cell wall exoskeleton was reduced in both the rigid core (**Figure 2A**) and mobile fractions (**Supplementary Fig. 9-10**). These results suggest that the outer eumelanin layer functionally compensates for the relatively hydrophilic inner wall composition of resting conidia and that loss of melanin triggers compensatory reinforcement of the polysaccharide scaffold.

**Figure 2.**
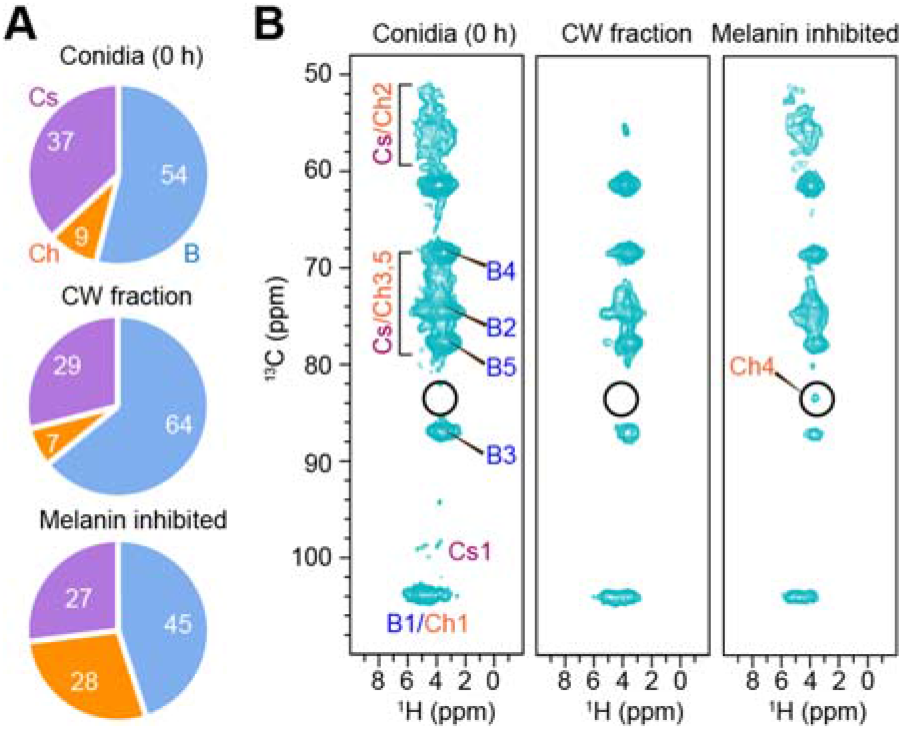
Melanin inhibition enhances chitin synthesis. (**A**) Composition of polysaccharides in untreated conidia, the cell wall fraction isolated after boiling SDS/DTT treatment, and melanin-inhibited conidia treated with BCS, quantified from deconvolution of 1D ^13^C CP spectra. (**B**) 2D ^13^C-^1^H hCH spectra of untreated conidia (left), cell wall fraction (middle), and melanin-inhibited conidia (right). Circles show the expected position for chitin ^13^C/^1^H4 peaks.

### Germ tube formation halts β-1,3-glucan synthesis and favors chitosan/chitin production

The early stages of germination are characterized by conidial swelling, which, surprisingly in *R. delemar*, occurred without disrupting the melanin layer. TEM revealed that swollen conidia retained a well-defined electron-dense outer wall, thinner to that of resting conidia (**Figure 3A**). These weakened sites preceded germ tube emergence and progressed into clear wall discontinuities during polarization (**Figure 3A**; right). Notably, the germ tube cell wall appeared as a single thinner layer derived from a newly synthesized inner wall originally beneath the plasma membrane in the conidia, highlighted by yellow arrowheads. This inner layer became progressively more prominent at the germination site and contributes to the newly synthesized cell wall of the growing hypha. The reduction in cell wall thickness (**Figure 3B**) occurred alongside a two-fold increase in conidial diameter during swelling (**Supplementary Fig. 11**). The thinner germ tube wall reflects ultrastructural reorganization and loss of the compact outer layers initially present in conidia. Taken together, these observations show that germ tube formation depends on *de novo* synthesis of a specialized cell wall architecture distinct from that of the resting conidium, likely due to the activation of different molecular pathways.

**Figure 3.**
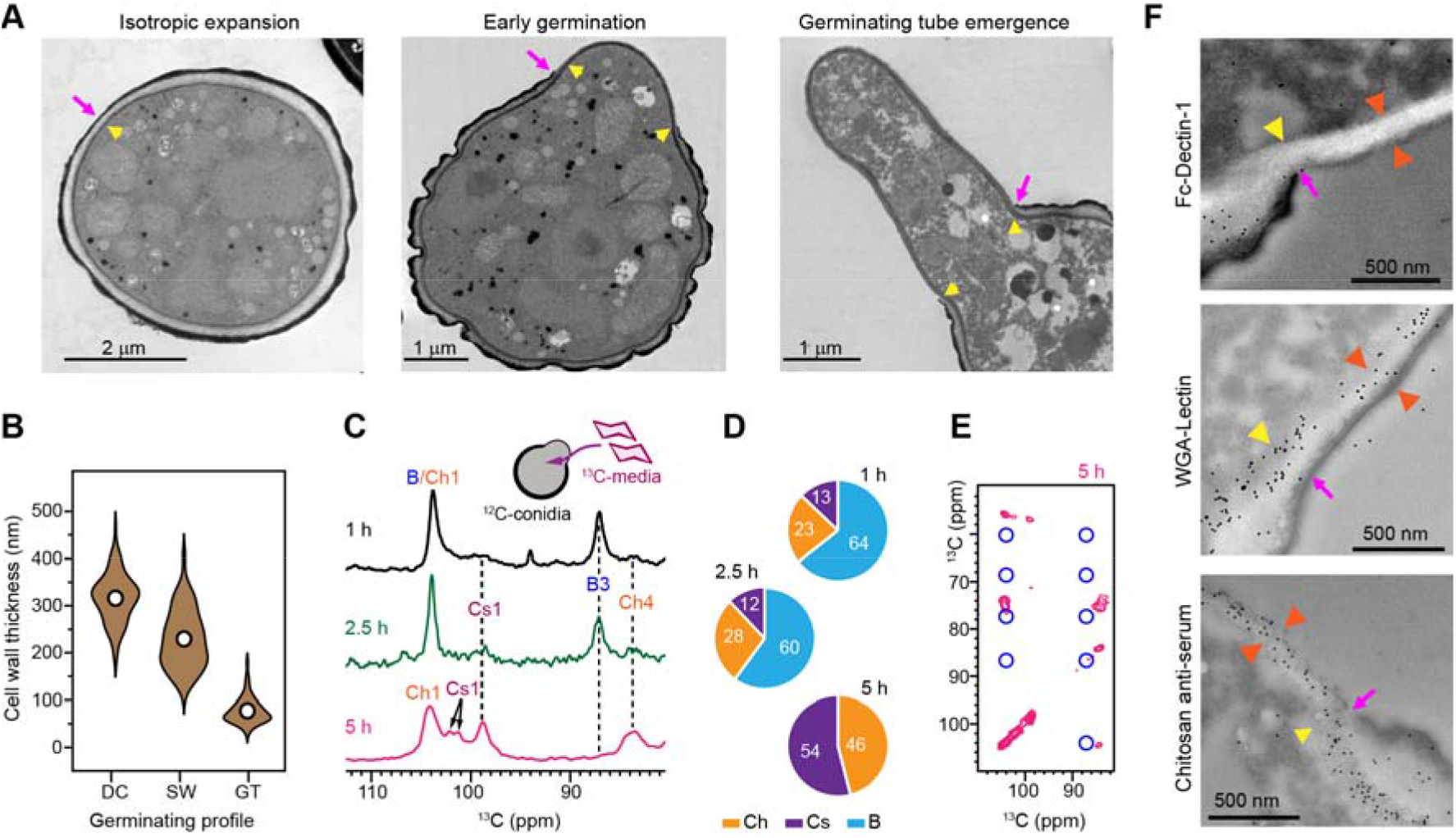
Time-resolved neosynthesis of cell wall polysaccharides during germination. (**A**) TEM images of germinating conidia showing swollen conidia undergoing isotropic expansion at 1 h (left), swollen conidia with rupture of the outer cell wall enabling emergence of a polarized germ tube (middle), and elongated germ tubes (right). Magenta arrows indicate changes in the outer wall, while yellow arrows highlight the thinner inner layer that extends and contributes to germ tube wall formation. (**B**) Violin plot showing the reduction in cell wall thickness from dormant conidia (DC) to swollen (SW) and germ tube (GT) stages (n=120 per group). (**C**) 1D ^13^C CP spectra detecting only neosynthesized polysaccharides in germinating ^12^C-conidia grown in ^13^C-labeled media (inset). Spectra were collected at different time points of 1 h (black), 2.5 h (green), and 5 h (magenta). Dashed lines highlight the signature peaks of chitin, β-1,3-glucan, and chitosan. (**D**) Molar composition of rigid neo-synthesized polysaccharides (%) in the 1-5 h culture. Compositional percentages are calculated by deconvoluting 1D ^13^C CP spectra. (**E**) 2D ^13^C-^13^C CORD spectra of the 5 h culture, with blue circles indicating the absence of β-1,3-glucan signals. (**F**) Cytochemical labeling of gold-specific staining for β-1,3-glucan (with Fc-Dectin-1, top), chitin (with WGA-lectin, middle), and anti-chitosan rabbit antiserum (bottom). Scale bars: 500 nm. Electron-dense gold particles (black dots) mark probe binding. The newly formed thin cell wall of germ tube (orange arrowheads) is labeled by WGA and anti-chitosan. Dectin labeling is only positive in the cell wall of the original resting conidium cell wall. Magenta arrows indicate the location of outer wall changes, and yellow arrows highlight the extending inner layer contributing to germ tube formation.

To track cell wall synthesis during the morphotype transition, unlabeled dormant conidia were grown in a ^13^C-labeled medium, such that only newly synthesized molecules incorporated ^13^C and were detected by ssNMR (**Figure 3C**). During the resting and swollen stages (2.5 hours), the neo-synthesized polysaccharides were primarily β-1,3-glucan, indicated by its carbon-3 (B3) peak at 87 ppm, along with minor contributions from chitosan, detected by the broad carbon-1 (Cs1) peak at 100 ppm, and chitin, detected by the carbon-4 (Ch4) peak at 83.5 ppm. By 5 hours, however, the β-1,3-glucan (B3) signal disappeared completely from the 1D ^13^C CP spectrum, while strong signals for chitosan (Cs1, 97-102 ppm) and chitin (Ch4, 83.5 ppm) emerged as the dominant features with a 54:46 ratio of chitosan to chitin in neo-synthesized cell walls (**Figure 3D**; **Supplementary Figs. 12**-**14**). The lack of β-1,3-glucan among newly synthesized rigid carbohydrates was validated using 2D ^13^C-^13^C correlation spectra, which provide enhanced resolution yet failed to reveal any characteristic peaks of this molecule (**Figure 3E**). Consistently, β-1,3-glucan signals were also absent in the mobile phase, confirming the complete halt of its biosynthesis (**Supplementary Fig. 15**).

Cytochemical analysis confirmed the polymer organization of the cell wall during conidial germination (**Figure 3F**; **Supplementary Fig. 16**). The thinner and extracellular regions of the germ tube wall were not labeled by Fc-Dectin-1, consistent with ssNMR data showing that, unlike resting conidia, β-1,3-glucan was largely absent from the germ tube cell wall. In contrast, the germ tube wall showed strong and uniform labeling with both WGA-colloidal gold and anti-chitosan antibody, confirming the presence of chitin and chitosan distributed throughout the wall without distinct subcellular localization.

These data reveal that cell wall expansion during conidial swelling largely preserves the pre-existing β-1,3-glucan-rich scaffold, whereas the transition to polarized growth involves extensive remodeling marked by β-1,3-glucan depletion and enrichment of chitin and chitosan required for hyphal tube formation. This shift reflects dynamic cell wall reorganization through coordinated polymer turnover and *de novo* biosynthesis.

### Growth polarization progressively and comprehensively remodels *R. delemar* cell walls

To distinguish synthesis, turnover, and reorganization of polysaccharides during germination, we combined three complementary ^13^C-labeling strategies (**Supplementary Fig. 17**). First, unlabeled conidia grown in ^13^C-labeled medium allowed selective detection of *de novo* polymer synthesis (**Figure 3C**): β-1,3-glucan was the major newly synthesized polysaccharide during early germ tube emergence, indicating strong glucan synthase activity, but its relative abundance decreased during later hyphal development, when chitin synthase and deacetylase activities dominated. Second, ^13^C-conidia grown in unlabeled medium only monitor pre-existing wall components, where loss of signal reflected polymer degradation by glycosyl hydrolases (**Supplementary Fig. 18**). Here, β-1,3-glucan was slightly reduced, chitin increased, and chitosan remained relatively stable during germination. Finally, ^13^C-conidia germinated in ^13^C-medium provided a comprehensive view of total cell wall architecture: the rigid wall remained compositionally stable during the resting and swollen conidial stages, whereas β-1,3-glucan decreased markedly relative to chitin and chitosan upon germ tube emergence (**Figure 4A**). Therefore, germination involves coordinated β-1,3-glucan turnover and progressive enrichment of chitin and chitosan through both polymer synthesis and hydrolysis, maintaining wall flexibility while supporting polarized hyphal growth.

**Figure 4.**
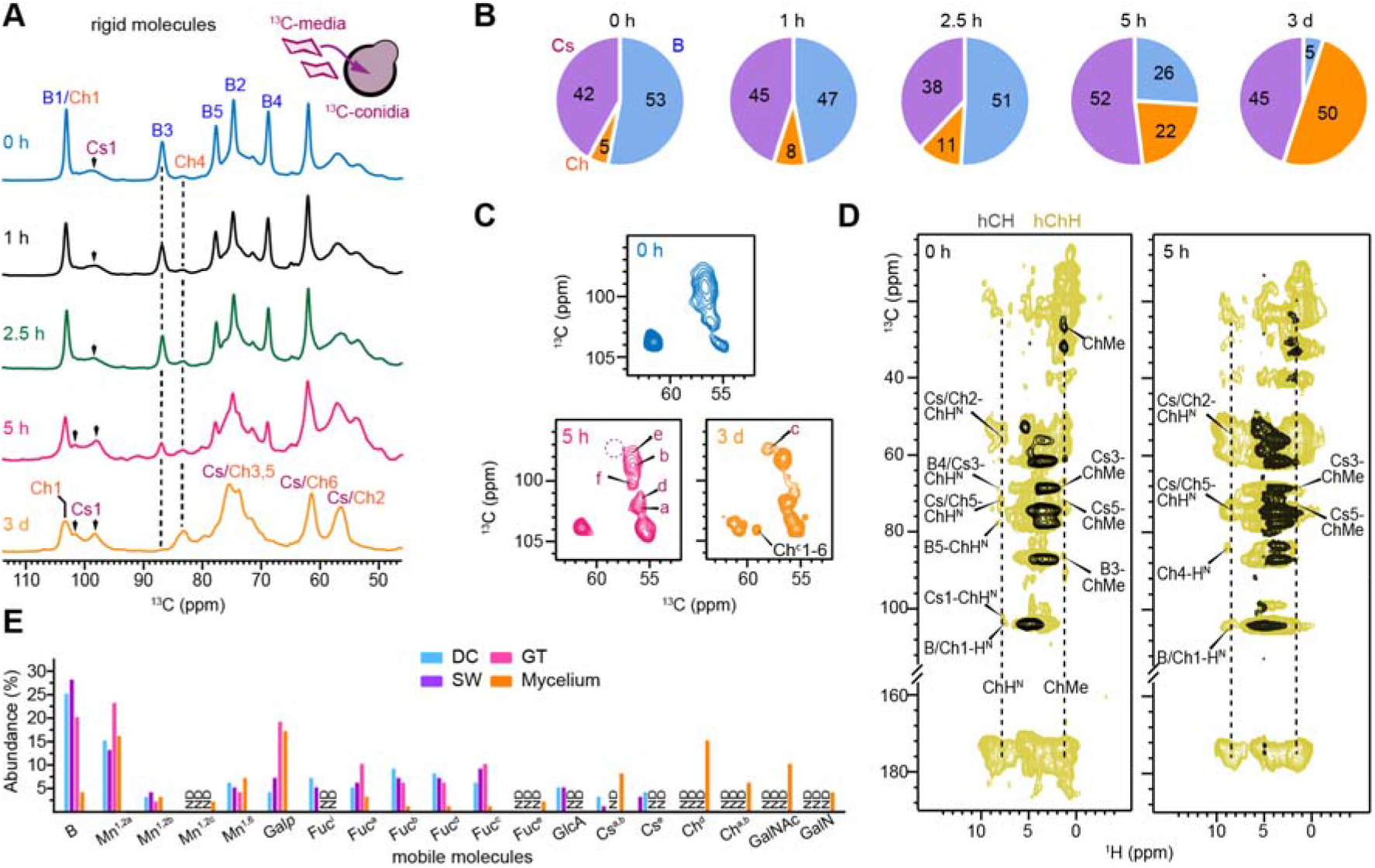
Progressive changes in carbohydrate structure, composition and interactions during germination. (**A**) 1D ^13^C CP spectra showing rigid cell wall polysaccharides at different developmental stages. The inset illustrates the labeling scheme in which ^13^C-labeled conidia were grown in ^13^C-labeled media. Samples correspond to 0 h and 1 h (resting conidia), 2.5 h (swollen conidia), 5 h (germ tube emergence), and 3 days (mycelium). Dashed lines indicate β-1,3-glucan carbon-3 (B3) and chitin carbon-4 (Ch4). Arrows highlight chitosan carbon-1 peaks, which shift from 2.5 h to 5 h during germ tube formation. In 3-day-old hyphae, the rigid fraction is dominated by chitin and chitosan. (**B**) Pie-charts showing the molar composition of rigid polysaccharides in *R. delemar* cell walls, quantified by deconvolution of 1D ^13^C CP spectra. (**C**) Zoomed view of C1 signals from 2D ^13^C CORD spectra showing distinct chitosan forms observed in cultures at 0 h (blue), 5 h (magenta), and 3 days (orange). (**D**) Overlay of 2D hCH spectra (black) showing polysaccharide signals and hChH spectra (pale yellow) measured with 0.5 ms RFDR mixing to detect long-range correlations. Data are shown for *R. delemar* conidia at 0 h (left) and after 5 h of germination (right). (**E**) Molar composition of mobile components at different culture germination stages calculated from intensity analysis of 2D ^13^C-DP refocused INADEQAUTE spectra. ND: not detected.

Compositional analysis of the rigid polysaccharides quantified the progressive depletion of β-1,3-glucan during germination (**Figure 4B**; **Supplementary Fig. 19**). β-1,3-glucan accounted for half of the rigid wall fraction in 0-2.5-hour cultures, decreased to a-quarter at 5 hours, and reached only 5% in the 3-day-old hyphal sample^14^. In contrast, chitin increased to 22% at 5 hours, coinciding with germ tube emergence, and reached 50% in mature hyphae. Because newly formed germ tubes lack detectable β-1,3-glucan, the residual signal at 5 hours likely originates from dormant and swollen conidial regions.

In addition to these quantitative changes, distinct morphotypes have been identified during germination. Upon germ tube emergence, the chitosan C1 (Cs1) region displayed two major peaks at 98 ppm and 102 ppm, in contrast to the single broad peak at 99 ppm observed in dormant and swollen conidia (**Figure 4A**), resembling the profile of mature hyphae and indicating that newly synthesized chitosan adopts a hypha-like structure. Higher-resolution view by 2D ^13^C correlation spectra showed that, although all five major chitosan forms were present throughout development (0-5 hours), germination was marked by selective enrichment of type-a and type-b chitosan, whereas type-f and type-d were substantially reduced (**Figure 4C**). A similar distribution was observed in mature hyphae, where type-a and type-b accounted for nearly two-thirds of total chitosan, indicating that type-f and type-d are characteristic of dormant and swollen conidia, whereas type-a and type-b are specifically associated with germ tube formation and polarized hyphal growth.

In later stages, β-1,3-glucan biosynthesis resumes at a low level to support formation of the mature hyphal network. This is supported by two observations. First, β-1,3-glucan still accounts for 5% of total polysaccharides in the 3-day-old hyphal sample (**Figure 4B**). Second, type-c chitosan, which is detected in hyphal cells, is absent from both conidial cells and germ tubes (**Figure 4C**). Recent work showed that β-1,3-glucan, type-c chitosan, and type-c chitin form a covalently linked carbohydrate complex removable by the chitin synthase inhibitor nikkomycin, indicating that this structure is unique to the mature hyphal network^14^.

During germination, the mechanical core of the cell wall was remodeled from a β-1,3-glucan-chitin scaffold into a binary rigid network primarily composed of chitin and chitosan. 2D ^13^C-^1^H correlation spectra revealed that dormant conidia contained weak intermolecular contacts between β-1,3-glucan and chitin on the sub-nanometer scale, including those between β-1,3-glucan carbon-5 and the chitin amide proton (B5-ChH^N^) and between β-1,3-glucan carbon-3 and the chitin methyl group (B3-ChMe; **Figure 4D**). These weak interactions disappeared during germ tube emergence due to the depletion of β-1,3-glucan. In parallel, new contacts formed between chitin and chitosan, including Cs5-ChMe and Cs3-ChH^N^, demonstrating structural reorganization of the wall toward a chitin-chitosan-dominated architecture.

The mobile wall fraction also underwent gradual compositional changes during germination but less prominent than the rigid polysaccharides (**Figure 4E**). Mannan and polymeric fucose persisted throughout this period, although their relative abundances fluctuated and specific structural allomorphs appeared or disappeared over time. Mobile β-1,3-glucan increased during conidial swelling but decreased markedly upon germ tube emergence (**Figure 4E**; **Supplementary Fig. 20**), consistent with the shutdown of β-1,3-glucan synthesis and increased chitin synthase and deacetylase activity during polarized growth (**Figures 3C, D**). The GlcA signals disappeared after the swelling stage, indicating that this polymer is primarily required in resting and early swollen conidia. Three chitosan forms were also detected in the mobile phase during early germination but were lost thereafter, suggesting their progressive incorporation into the rigid wall through stronger association with chitin.

As development progressed toward the mycelia, the mobile fraction became increasingly enriched in chitin, chitosan, and other cationic polysaccharide units. This shift from a chemically diverse mobile matrix in germ tubes toward one dominated by structurally reinforcing polymers in mature mycelia indicates progressive wall consolidation and suggests the formation of a more interactive and mechanically robust matrix that supports mature hyphal growth and environmental resilience.

### Neutrophils bind and phagocytose early germinated conidia of *R. delemar*

As β-1,3-glucan, chitin, and chitosan are fungal PAMPs, their stage-specific remodeling during germination is expected to alter host immune recognition. We therefore examined how human neutrophils (NIR^+^, blue mask analysis) recognize resting, swollen, and germinating conidia of *R. delemar* (Green^+^, pink fungal mask analysis) using live-cell imaging (**Supplementary Fig. 21**)^26,28^. Image segmentation of fungal and neutrophil masks enabled the identification and quantification of individual cells and their interactions with *R. delemar* (Green^+^/NIR^+^, red mask analysis, **Figure 5A, B**).

Analysis demonstrated that dormant conidia (0 h) came into contact with neutrophils and were phagocytosed (**Figure 5C**). After 2 h of fungal growth, neutrophils exhibited rapid recruitment to swollen conidia, with detectable binding occurring between 15 and 60 min (**Figure 5A**; **Supplementary Fig. 21**). Once a neutrophil bound to *R. delemar*, it attracted nearby neutrophils, resulting in cluster formation. Comparable binding kinetics were observed after 4 h of fungal growth, including binding to the surface of conidial ghost (orange arrowhead) and germ tubes (blue arrowhead; **Figure 5B**).

**Figure 5.**
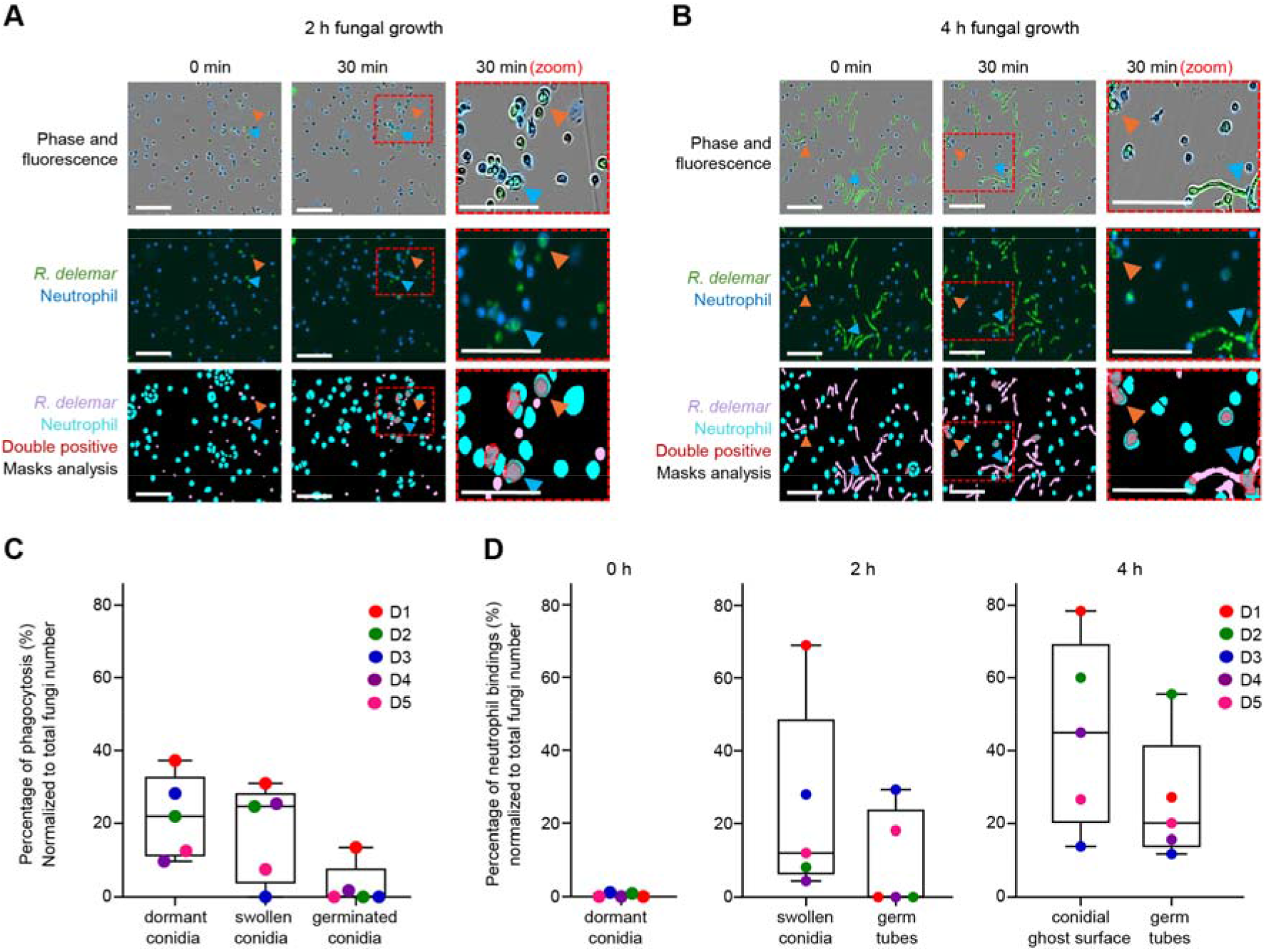
Neutrophils bind avidly to germinating conidia of *R. delemar*. (**A**) Images showing neutrophil binding to *R. delemar* at 2 h growth. Human neutrophils were incubated with *R. delemar* resting conidia or with fungi pre-grown in RPMI medium for 2 h prior to co-culture. Neutrophils were labeled with a near-infrared (NIR) tracker, and *R. delemar* was stained with Sytotox Green. Live-cell imaging was performed for with images acquired at 30 min to monitor neutrophil-fungus interactions. Image segmentation (mask analysis) identified *R. delemar* (pink), neutrophils (blue), and interaction sites (red). Orange arrowheads identify conidial surface, and blue arrowheads identify germ tubes/hyphae. Scale bar: 100 μm. (**B**) Image showing neutrophil binding to *R. delemar* at 4 h growth. Live-cell imaging was performed with images acquired at 30 min. (**C**) Quantification of double-positive (NIR□/green□) objects at 0-4 h growth of *R. delemar* phagocytosed by neutrophils. Data is from n = 5 independent donors (D1-5) and are shown mean ± s.e.m. (**D**) Quantification of double-positive (NIR□/green□) objects at 0-4 h growth of *R. delemar* across resting conidia and swollen conidia and germinating conidia. Data is from n = 5 independent donors (D1-5) and are shown mean ± s.e.m.

Quantification of neutrophil-fungus interactions was performed by measuring double-positive objects (NIR□/green□, red mask analysis, **Figures 5A, B**) and determined specific fungal target regions defined by conidial surface and germ tubes. Neutrophils binding affinity was highest for swollen conidia (2 h; **Figure 5D**). The total number of neutrophil binding events gradually increased, indicating that fungal surface expansion increases the likelihood of neutrophil attachment (**Supplementary Fig. 21**). After 4 h of fungal growth, approximately 40% of neutrophil binding events occurred at the conidial ghost surface, whereas around 20% targeted the germ tubes, with a preference for binding to the apical zones of the germ tubes, indicating that expanding *R. delemar* germ tubes continuously attract neutrophils (**Figure 5D**).

Additionally, the swollen conidia stage of *R. delemar* enhanced the capacity of neutrophils to phagocytose conidia; however, this capacity decreased once conidia began to form germ tubes (**Figure 5C**). To determine whether the absence of β-1,3-glucan in germinating conidia depends on growth conditions, parallel cultures were grown in RPMI and minimal medium under both CO_2_-enriched and standard conditions. Fc-Dectin-1 labeling yielded consistent results across all conditions, with no detectable β-1,3-glucan in germ tubes (**Supplementary Fig. 22**), confirming that this feature is conserved during germination and is independent of growth conditions. Accordingly, the absence of β-1,3-glucan as a PAMP does not diminish the neutrophil response. This proves that stage-specific cell wall remodeling during germination alters outer wall components, and that their exposure to immune cells likely shapes neutrophil recognition, potentially enhancing pathogenic success through morphotype heterogeneity.

## DISCUSSION

This study provides a molecular-level framework for understanding cell wall remodeling during fungal germination in *R. delemar* and reveals a developmentally programmed transition from a β-1,3-glucan-rich dormant architecture to a chitin-chitosan-dominated polarized wall. Integrating multidimensional ssNMR with ultrastructural analysis, we show that resting conidia possess a rigid core enriched in β-1,3-glucan, with contributions from multiple polymorphic forms of chitosan and a minor fraction of chitin, embedded within a chemically diverse mobile matrix containing mannans, fucose-rich polymers and glucuronic acid. The presence of specific polymers in both rigid and mobile phases highlights a continuum of structural organization rather than discrete stratification, suggesting that dynamic flexibility is intrinsically encoded within the conidial wall. These findings further confirm that the cell wall of Mucorales differs fundamentally from that of filamentous ascomycetes and basidiomycetes^29,30^.

A central finding of this work is the disappearance of rigid β-1,3-glucan during germ tube emergence and its replacement by a chitin-chitosan-dominated scaffold that supports polarized growth. Germination therefore does not represent simple wall expansion, but a coordinated remodeling process involving β-1,3-glucan synthase cessation, increased glycosyl hydrolase activity, enhanced chitin synthase and deacetylase activity, and selective enrichment of specific chitosan polymorphs. The progressive formation of direct chitin-chitosan intermolecular contacts further demonstrates that the mechanical core of the wall is reorganized from a β-1,3-glucan-containing conidial scaffold into a binary rigid network optimized for hyphal extension. This developmental switch observed in *Rhizopus* contrasts with canonical models in ascomycetes such as *Aspergillus fumigatus*, where β-1,3-glucan synthesis remains prominent during hyphal growth, and suggests that mucoralean fungi rely primarily on a chitin-chitosan scaffold for mechanical stability during directional growth.

The close morphological and genetical relationship between *R. delemar* and *Rhizopus oryzae* and, together with the published genome of *R. oryzae*, provides a framework for interpreting these structural changes^29^. *R. oryzae* has a large repertoire of carbohydrate-active enzymes (CAZymes) involved in cell wall synthesis, remodeling, and hydrolysis, including enzymes for chitin modification and recycling (34 chitin deacetylases), chitosan metabolism (43 enzymes including 3 chitosanases), and β-1,3-glucan synthesis (27 enzymes including 3 putative β-1,3-glucan synthases)^29,31^. As shown in model fungi such as *Saccharomyces cerevisiae* and *A. fumigatus*, enzymes within the same CAZyme family often perform distinct and specialized rather than redundant functions^32,33^. Similar functional specialization may explain the developmental specificity of polysaccharide remodeling in Mucorales and may also contribute to the limited efficacy of carbohydrate-active inhibitors targeting chitin and glucan synthesis. One notable example is that caspofungin, which inhibits β-1,3-glucan synthase, reduces brain fungal burden, and improves survival in murine disseminated mucormycosis at low, but not high, doses^34^. Unfortunately, gene deletion approaches have just barely started with this fungus and other Zygomycetes, making it impossible to understand their function as it has been done for the CAZYmes of ascomycetous fungi^35-37^.

The observed structural changes are also likely to influence host-pathogen interactions. Upon inhalation of conidia from *Rhizopus* and other zygomycetes, the host mounts a strong innate immune response in which fungal cell wall components play a central role in the activation of this response^38^. The heterogeneous surface exposure of β-1,3-glucan in dormant and germinating conidia, together with the presence of chitosan, which has recognized immunological functions during infections caused by filamentous fungi such as *Aspergillus*^39^, may contribute to disease establishment, as β-1,3-glucan is a major PAMP recognized by host receptors including Fc-Dectin-1 and complement receptors. Although chitin is generally considered immunologically inert, it can also participate in host immune responses, and the incorporation of glucosamine residues into the linear chitin chain may further alter host recognition of this modified polysaccharide^40-42^.

The persistence of a melanin and protein rich outer layer during conidial swelling suggests that these outer wall components contribute to the hydrophilicity of *Rhizopus* conidia. However, the specific cell wall components responsible for wall permeability in mucoralean conidia remain unknown. The conidial outer layer differs markedly between ascomycetes and *Rhizopus* conidia; notably, the absence of rodlet hydrophobins on the surface of *R. delemar* may be one reason for the hydrophilic nature of zygomycete conidia^43^. To date, no proteomic studies have investigated the composition of surface cell wall proteins or their role in protection against reactive oxidants. Our observations further indicate that the melanin layer in *R. delemar* is highly elastic and water permeable, distinguishing it from the external pigment layer of ascomycetous filamentous fungi such as *Aspergillus*, where germination is consistently associated with melanin layer breakdown^44^. This study also suggests a potential association between chitin and melanin, similar to that described in *Cryptococcus*, where chitin acts as a scaffold for the progressive incorporation of melanin pigments into the cell wall^45^. Alternatively, melanin could be anchored to β-glucans in *R. delemar*, as the highest abundance of β-1,3-glucans is detected in melanin-associated conidia; however, this possibility remains unexplored. Although the role of tyrosinase/laccase in conidial melanin synthesis has been supported by the use of inhibitors, the complete biosynthetic pathway remains unresolved both biochemically and genetically^46^.

The limited compositional changes and persistence of mobile polysaccharides in both resting conidia and germ tubes suggest that they are unlikely to serve a primary structural role; instead, they may unspecifically fill the joints between cell-wall structural bricks, contribute to hydration control, or mediate extracellular interactions. The origin of GlcA remains unknown and may derive from glycoproteins or from GlcA-containing fucans. Because fucans are present in high abundance, they may contribute to the immunological or inflammatory response to *Rhizopus* during infection, although this possibility has not been investigated. Whether *Rhizopu*s fucans perform functions analogous to the sulfated fucoidans of algae and echinoderms, despite lacking sulfation, remains unknown. In plants, fucans play important roles in defense against external microbial aggressors, raising the possibility that fungal fucans may also have specialized protective or immunomodulatory functions^47,48^. The mobile matrix also shows morphotype-specific regulation, with GlcA restricted to dormant and swollen conidia and mobile chitosan gradually incorporated into the rigid scaffold through germination.

Collectively, these findings shift the paradigm of mucoralean cell wall architecture by showing that polarized growth is driven by coordinated polymer turnover, selective biosynthetic shutdown, polymer redistribution, and altered intermolecular association patterns. This work establishes a molecular framework linking morphogenesis, mechanical adaptation, and immune modulation during germination of *R. delemar*. More broadly, it highlights fundamental differences between early-diverging fungi and canonical fungal models and demonstrates the power of ssNMR spectroscopy to resolve dynamic cell wall organization at molecular resolution.

## METHODS

### Preparation of resting and germinating conidia for ssNMR

Conidia of *R. delemar* (FGSC-9543) were produced on solid medium with 1.5% agar containing 10 g/L dextrose, 2.5 g/L ammonium sulfate, and 1.7 g/L yeast nitrogen base without amino acids and ammonium sulfate (DF0335-15G, Fisher Scientific), and incubated for 3 days at 37 °C. For the germination assays, the same medium composition was used without agar. For preparation of isotopically labeled conidia and germination in labeled growth medium, ^13^C-glucose (CLM-1396-PK, Cambridge Isotope Laboratories) and ^15^N-ammonium sulfate (NLM-713-PK, Cambridge Isotope Laboratories) were used in place of natural-abundance substrates. The medium pH was adjusted to 6.5 and sterilized by autoclaving.

After sterilization, the medium was poured into Petri dishes and allowed to solidify under a sterile laminar flow hood. Conidia were harvested by suspending the cultures in 0.5% Tween solution. The conidial suspension was filtered through muslin cloth to remove mycelial fragments. The filtrates were examined under a light microscope to confirm the absence of mycelial contamination.

Conidia were quantified using a hemocytometer, and 10^6^-10^8^ conidia cells were incubated into 40 mL of liquid medium in a 100 mL Erlenmeyer flask at 37°C and 250 rpm for up to 5 h. The fungal morphologies, including resting conidia, swollen conidia (diameter of the conidium <1.8 × that of resting conidia), and germ tubes (germ tubes > 0.9 × conidial diameter) were quantified at defined germination time points (1 h, 2.5 h, 4 h, and 5 h; **Supplementary Table 1**).

Three different experimental conditions were used to investigate cell wall remodeling during germination. In the first condition, unlabeled resting conidia were germinated in ^13^C-labeled medium (unLCLM) to track newly synthesized components, In the second condition, ^13^C labeled conidia were germinated in unlabeled medium (LCunLM) to monitor the fate of preexisting cell wall material. In the third condition, ^13^C-labeled conidia were germinated in ^13^C-labeled medium (LCLM), allowing for comprehensive observation of both pre-existing and newly synthesized polysaccharides.

### TEM analysis of cell wall morphology

For TEM observations, the fungal samples were fixed in 2.5% glutaraldehyde and 2% paraformaldehyde (15700, Electron Microscopy Sciences) in 0.1□M cacodylate buffer,followed by embedding in 2% agarose and post-fixation in 0.1□ M osmium tetroxide (SKU 19152, Electron Microscopy Science). Dehydration was achieved through a graded acetone series of increasing concentrations, followed by infiltration with epoxy resin and acetone at ratios of 25:75, 50:50, and 75:25 (v/v), respectively. Samples were incubated in the 75:25 epoxy resin-acetone solution overnight and then treated with 100% resin for two days with multiple resin changes. Finally, the sample was placed in an oven at 70□°C to prepare the blocks. Ultrathin sections were stained with 1% uranyl acetate (22400, Electron Microscopy Sciences) and lead acetate (17800, Electron Microscopy Sciences) and mounted on carbon-coated grids (FCF-150-CU, Electron Microscopy Sciences). TEM imaging focused on sample cross-sections, and 100 cell wall thickness measurements were performed for each group (**Supplementary Table 1**). Cell wall thickness was measured using ImageJ software, and the data were analyzed statistically using a t-test (**Supplementary Table 1**).

### Cytochemical labeling

Fungal cells were fixed in 4% paraformaldehyde and 0.5% glutaraldehyde prepared in PBS. Following fixation, samples were rinsed in PBS, dehydrated through a graded ethanol series, and infiltrated with LR White resin. Embedding was performed under oxygen-free conditions, and polymerization was carried out by oven curing for 24 h. Ultrathin sections were prepared using an Enuity Leica ultramicrotome (Leica Microsystems, Austria) and collected on 200 mesh formvar-carbon gold grids for immunogold labeling. All grids were observed under a JEOL JEM 1400 TEM at 20 kV at the Center for Advanced Microscopy of Michigan State University (MSU).

For β-1,3-glucan localization, grids were subjected to aldehyde neutralization by incubation in 0.2 M glycine in PBS (pH 7.4) for 45 min to quench residual reactive aldehydes. Grids were washed with endotoxin-free water and incubated for 1 h with 100 µg mL^-1^ recombinant human Fc-Dectin-1 (Fc-hDectin-1a, InvivoGen). After washing, grids were incubated for 1 h with a 1:10 dilution of Protein A-colloidal gold conjugate (10 nm, Aurion) in PBS containing 1% bovine serum albumin (BSA). Grids were washed three times with PBS and three times with HPLC-grade water prior to imaging. For specificity controls, the same labeling procedure was performed with one modification: Fc-hDectin-1 was preincubated with β-1,3-glucan (CAS # sc-213514, Santa Cruz Biotechnology, Inc.) at 1:1 ratio for 1 h before application to the grides, followed by the same steps described above.

Chitin localization was performed using wheat germ agglutinin (WGA) conjugated to 20 nm colloidal gold particles (*Triticum vulgaris* lectin). Prior to labeling, samples were incubated in 0.2 M glycine in PBS for 45 min, followed by washing with PBS supplemented with 1% bovine serum albumin (BSA). Grids were then incubated with the 1:9 dilution of WGA-gold conjugate solution with PBS-1% BSA for about 1 h under gentle mixing. After labeling, grids were washed with PBS and rinsed with HPLC-grade water for TEM imaging. For control experiments, a solution containing WGA–lectin gold conjugate (EY, CAS GP-2102-20; 1:10 dilution), chitin oligosaccharide, and the chitin-binding inhibitor N,N’,N’’-triacetylchitotriose (50 mM; Fisher, CAS#T2144) was prepared. Chitin was added at final concentrations corresponding to 1:100 and 1:1000 dilutions to block WGA binding. Grids were first incubated in 0.2 M glycine in PBS for 45 min, followed by washing with PBS containing 1% bovine serum albumin (BSA). The grids were then incubated in the control solution for 1 h. After labeling, grids were washed with PBS and rinsed with HPLC-grade water prior to TEM imaging.

Chitosan-specific rabbit antiserum and control pre-immune serum were obtained from Eurogentec. Immunization was performed with Chitosan low molecular weight (448869>80% glucosamine)1 mg of chitosan with complete Freund’s adjuvant, followed by booster injections every two weeks with 0.5 mg of chitosan in incomplete Freund’s adjuvant. Serum specificity was tested by ELISA using anti-chitosan antiserum and ELISA. Pre-immune sera were used as a control Rabbit sera were diluted to 1:100 in PBS containing 1% BSA and filtered through a 0.22 μm syringe filter. After incubation, grids were washed with PBS containing 1% BSA and then incubated with a 1:10 dilution of Protein A-Gold conjugate (Aurion) for 1 h, followed by washing with PBS and then HPLC-grade water.

Mannan localization was performed using Concanavalin A lectin (ConA) (EY Laboratories, Inc, GP-1104-10) conjugated to 10 nm colloidal gold particles. Grids were incubated at a dilution of 1:10 gold conjugates in PBS supplemented with 1mM CaCl_2_, 1mM MnCl_2_, and 1% BSA. After labeling, grids were washed with PBS and rinsed with HPLC-grade water.

### Extraction of the cell wall with SDS/DTT for ssNMR

Labeled conidia (∼10^8^ spores/mL) were harvested from the agar plates as described above. The conidial suspension was transferred into a Teflon ball mill jar along with 0.5 mm glass beads at a 1:1 (v:v) ratio. The material was ball-milled in a Retsch MM200 mixer mill at 25 Hz for 2 min to disrupt the conidia, and the extent of cell wall breakage was assessed microscopically. The glass beads were removed from the sample by decantation. The resulting cell wall material was then boiled twice for 15 min in a mixture of 2% Sodium dodecyl sulfate (SDS)^49^, 40 mM dithiothreitol (DTT)^50^, and 50 mM Ethylenediaminetetraacetic acid (EDTA) to remove membrane-associated proteins and lipids^30,33^. The sample was then centrifuged to remove excess particles and packed into a 1.6 mm rotor for NMR analysis. The relative abundance of the carbohydrate components of the extracted cell wall was determined via deconvolution of 1D ^13^C CP spectra (**Supplementary Table 2**).

### Melanin inhibition

To inhibit melanin biosynthesis in conidia, a 10 mM stock solution of bathocuproinedisulfonic acid disodium salt (BCS) was prepared^51,52^, sterilized by autoclaving and stored at 4 °C until use. To prepare the BCS-supplemented medium, 1 g of ^13^C-glucose, 0.25 g of ^15^N-ammonium sulfate, 0.17 g of YNB, and 2% of agar were dissolved in 100 mL of Millipore water. The pH was adjusted to 6.5, and the medium was autoclaved. Once the autoclaved medium had cooled to approximately 60 °C, BCS was added to a final concentration of 1 mM and mixed thoroughly. The BCS-supplemented medium was inoculated, and the plates were incubated at 28 °C for 5 days. After incubation, the conidia (∼10^6^ spores/mL) were recovered in a Tween 20 solution, purified by Miracloth filtration, and packed into a 1.6 mm rotor for ssNMR analysis. The relative abundance of the carbohydrates in melanin-inhibited sample was determined by deconvoluting 1D ^13^C CP spectra (**Supplementary Table 2**).

### ^13^C Solid-state NMR analysis of cell wall composition and dynamics

SsNMR experiments were conducted on a Bruker Avance Neo spectrometer (18.8 Tesla) housed at MSU Max T. Rogers NMR facility. Most experiments were performed using a 3.2 mm HCN triple-resonance probe at magic-angle spinning (MAS) frequency of 15 kHz at 290 K. ^13^C chemical shifts were externally referenced to the tetramethylsilane (TMS) scale using the tripeptide N-formyl methionine-leucine-phenylalanine (f-MLF)^53^ and are listed in **Supplementary Table 3**. Typical radiofrequency (rf) field strength ranged from 71.4 to 83.3 kHz for ^1^H decoupling and hard pulses, while ^13^C hard pulses were applied at 50 kHz. Additional experimental parameters are provided in **Supplementary Table 4**.

Three types of 1D ^13^C spectra were acquired using varied polarization methods to differentiate molecular components based on their dynamics. Rigid components were identified using dipolar-coupling-based 1D ^13^C CP with a 1 ms contact time, maintaining Hartmann-Hahn matching conditions at 62.5 kHz for both ^13^C and ^1^H. The CP spectra were deconvoluted using the DMfit software to estimate the molar composition of rigid polysaccharides^54^. The results are summarized in **Supplementary Table 5**. Mobile molecules were examined using 1D ^13^C DP with a short recycle delay of 2 s, which preferentially detects molecules with rapid ^13^C-T_1_ relaxation. Quantitative detection of all molecules and carbon sites was achieved by applying longer recycle delays (35 s) to ensure complete relaxation to equilibrium before the acquisition of the next scan.

Resonance assignments were further confirmed by analyzing rigid components using 2D ^13^C-^13^C 53-ms CORD experiments^55^, which probe through-space correlations and were recorded for most samples. These experiments revealed intramolecular cross-peaks between different carbon sites in rigid molecules (**Supplementary Table 6**). To analyze mobile molecules, 2D ^13^C DP refocused J-INADEQUATE experiments were performed with a 1.5 s recycle delay (**Supplementary Table 7**)^56^. This method correlates a double-quantum (DQ) chemical shift with two corresponding single-quantum (SQ) chemical shifts in a covalently linked carbon pair, producing asymmetric spectra that helps track carbon connectivity through bond, thereby enabling identification of molecular backbones. Chemical shifts of rigid and mobile molecules are documented in **Supplementary Table 3**.

### Carbohydrate composition in *R. delemar* cell walls

Relative molar compositions of rigid and mobile carbohydrates were determined using complementary solid-state NMR approaches. The mobile fraction was quantified by peak-volume analysis of 2D ^13^C-DP refocused J-INADEQUATE spectra, whereas the rigid fraction was quantified using peak-volume analysis of 2D ^13^C-CP CORD spectra together with deconvolution of 1D ^13^C spectra. Peak volumes and intensities were extracted using Bruker TopSpin (version 4.1.4). To minimize uncertainties arising from spectral overlap, only well-resolved, unambiguous resonances were included in the analysis. For the 2D spectra, the volumes of assigned cross-peaks corresponding to individual polysaccharides were averaged. For the 1D spectra, deconvolution was performed within defined spectral regions, and the integrated peak areas were normalized to calculate the relative contribution of each component.

All spectra compared across samples were acquired and processed under identical experimental conditions. 1D peak areas and 2D peak volumes were normalized to the number of scans. Relative molar abundances were calculated by normalizing the integrated peak volumes or areas to the number of contributing resonances for each carbohydrate species and are reported as fractions of the total carbohydrate signal within the respective rigid or mobile domains. These values represent relative molecular distributions rather than absolute concentrations as summarized in **Supplementary Tables 2, 5**, and **8**. Standard errors were estimated from the variability of integrated peak volumes across selected resonances, and overall uncertainties were calculated by propagating individual errors.

### Fast MAS ^1^H-detection ssNMR for carbohydrate analysis

Proton-detection experiments, including 2D hCH and hChH, were acquired on two 800 MHz (18.8 Tesla) and 600 MHz (14.4 Tesla) Bruker Avance Neo spectrometers housed at MSU Max T. Rogers NMR facility^57,58^. For *Rhizopus* conidia, BCS-treated or melanin-inhibited sample, and cell wall fraction from SDS/DTT extraction, the ^1^H-detection ssNMR experiments were performed on the 800 MHz NMR equipped with a triple-resonance Phoenix 1.6 mm MAS probe at a MAS rate of 40 kHz. A small amount of sodium trimethylsilylpropanesulfonate (DSS) was added to the sample to reference the ^1^H chemical shifts. The ^13^C chemical shifts were externally referenced to the TMS scale using the adamantane methylene peak at 38.48 ppm. The rf field strength for the 90º pulse was set to 100 kHz for ^1^H and 50 kHz for ^13^C. The Hartmann-Hahn condition for CP was optimized to have a ^1^H rf field strength of 84 kHz, using a ramped amplitude (70%-100%), and a ^13^C rf field strength of 42 kHz. SPINAL-64 heteronuclear decoupling was applied on the ^1^H channel during acquisition. The CP contact time was optimized to 600 µs. For each ^1^H-^13^C CP spectrum, 1024 scans were co-added with a recycle delay of 3 s. The temperature of the cooling gas was set to 278K. The actual sample temperature was calculated from the difference in chemical shifts between the water and DSS peaks within each sample. The resulting temperatures were 284 K for the cell wall fraction, 292 K for the melanin-inhibited sample, and 298 K for the conidia samples. 2D data were collected using the States-TPPI method^59^.

For analysis of long-range intermolecular interactions, 2D spectra of *R. delemar* conidia and the 5 h germination time point were measured on the 600 MHz NMR at 60 kHz MAS. 2D hCH experiments were performed both without RFDR mixing for through-bond correlation analysis and with different second CP contact times to measure short- and long-range correlations. Short-contact hCH experiments used a second CP contact time of 50 µs, whereas long-contact experiments used a CP contact time of 500 µs. 2D hChH experiment employed a 0.8 ms RFDR-XY16 (radiofrequency-driven recoupling) mixing period^60,61^. For both hCH and hChH experiments, 192 time-domain points were acquired in the indirect dimension, with 512 transients accumulated per increment and a recycle delay of 2.0 s. Heteronuclear dipolar decoupling during the t_1_ evolution period was achieved using the slpTPPM (swept low-power two-pulse phase modulation) sequence^62^. During the direct ^1^H-detection period, WALTZ-16 decoupling was applied on the ^13^C channel with a rf field strength of 12.5 kHz. Water suppression was accomplished using the MISSISSIPPI sequence on the ^1^H channel^63^, with rf field strength of 15.2 kHz applied for 100 ms. The ^1^H and ^13^C chemical shifts of *R. delemar* polysaccharides, together with long-range intermolecular cross peaks and the experimental parameters, are reported in **Supplementary Table 9-11**.

### Binding of neutrophils to resting and germinating conidia

*R. delemar* was grown on Sabouraud dextrose agar slants for 5 days. Conidia were harvested in water containing 0.1% (v/v) Tween-20 and filtered through a 40 µm cell strainer. *In vitro* cultures were realized in RPMI 1640 medium (61870127, Gibco) containing 10% fetal calf serum (FCS) and 1% Penicillin-Streptomycin. Fungal growth of *R. delemar* conidia, deposited in wells of a 96-well plate, was continuously monitored at 37°C by an IncuCyte SX5 live-cell analysis system (Sartorius, Göttingen, Germany) for 7 h. Sytox Green (20 nM, S7020, Thermo Fisher Scientific) was added prior to neutrophil interaction and maintained throughout the entire experiment, staining *R. delemar*.

Neutrophils were isolated from blood of mixed healthy donors, by magnetic labeling (negative selection, MACSxpress whole blood neutrophil isolation kit, human, 130-104-434). Neutrophils were labelled with deep red CellTracker during 30 min before washing and the experiment (1μM, C34565, Thermo Fisher Scientific). The 0-4 h growth and binding affinity of 1×10^3^ *R. delemar* conidia and neutrophils at a ratio were monitored at 37°C by an IncuCyte SX5 live-cell analysis system every 5 min for 60 min. Hyphae length was analyzed using the NeutroTrack module of the Incucyte Analysis software. Image analysis of the binding of neutrophils to *R. delemar* was performed using the Incucyte Basic Analysis software. Neutrophils were detected in the near-infrared (NIR) channel and fungi in the green fluorescence channel. Double-positive objects, representing neutrophil-fungus interactions, were defined as overlapping NIR and green signals. Object counts of double-positive events were quantified to obtain the total number of interactions within the analyzed area. Neutrophil binding events to the fungi were determined after 60 min based on their different locations, such as the swollen conidia ghosts, germ tubes, and apical regions.

## DATA AVAILABILITY

All the original ssNMR data files will be deposited in the Zenodo repository, and the access code and DOI will be provided for public access.

## ACKNOWLEDGMENT

This project was supported by the National Institutes of Health (NIH) grant R01AI173270 to T.W. B.B. acknowledges funding from the Association Vaincre la Mucoviscidose/Association Grégory Lemarchal (RF20230503265 and RF20240503504) and the Agence Nationale de la Recherche (ANR-23-CE15-0013-03, ANR-24-CE15-2995-01, and ANR-25-CE15-1868-01).

## Notes

### Competing Interest Statement

The authors have declared no competing interest.

