## Supplementary file for "Dynamic Reprogramming of Fungal Cell Walls Underlies Germination and Immune Exposure in Zygomycetous Fungal Pathogens"

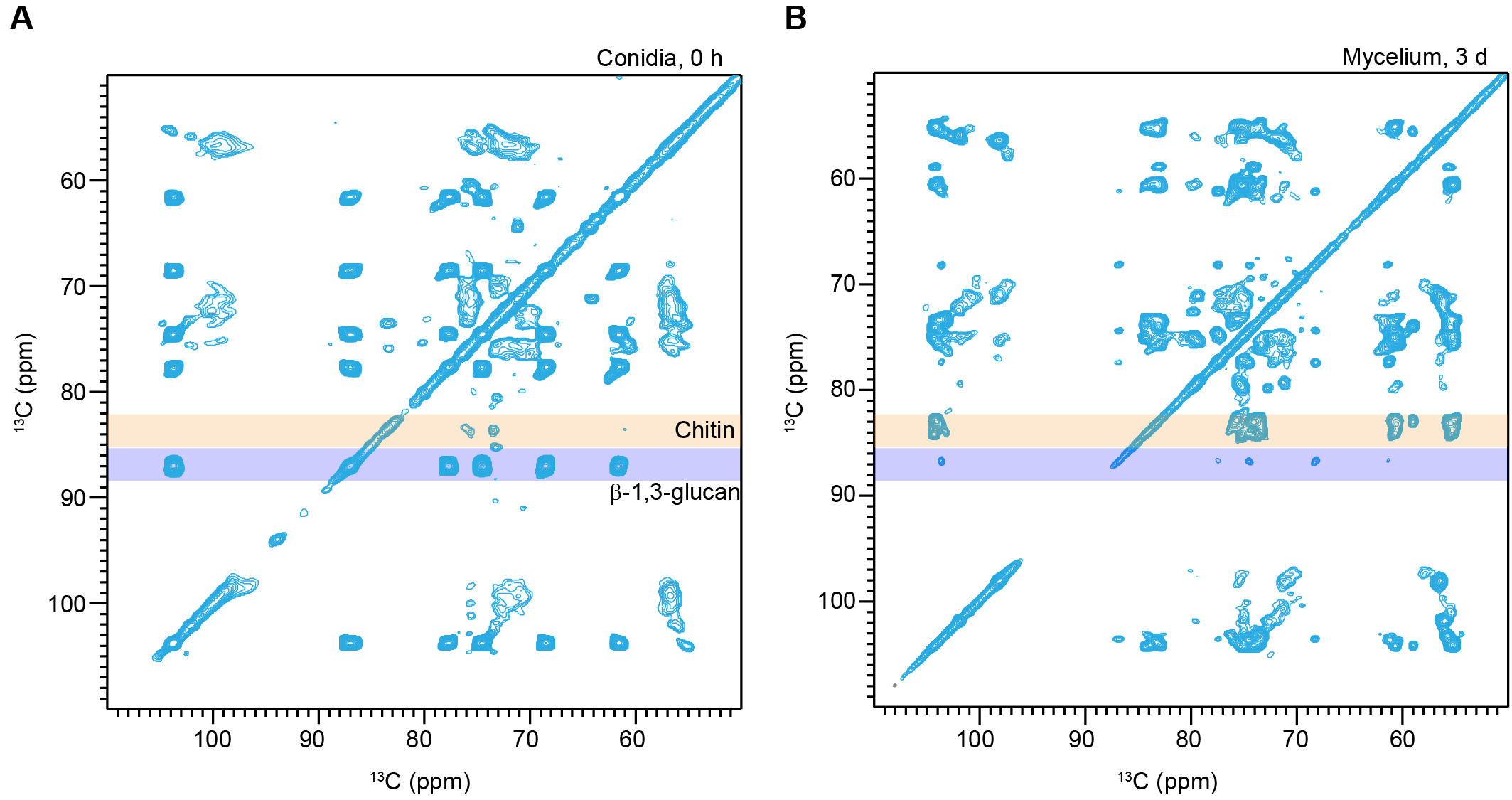

**Supplementary Figure 1.** **Carbohydrate signals of *R. delemar* conidia and mycelia**. (**A**) Carbohydrate region of 2D ^13^C-^13^C CORD spectrum for *R. delemar* conidia, representing the dominating signals for β-1,3-glucans (purple band at 87 ppm). (**B**) Carbohydrate region of 2D ^13^C-^13^C CORD spectrum for *R. delemar* mycelium shows a reduction in β-1,3-glucan signals, accompanied by a significant increase in chitin (orange band at 83 ppm).

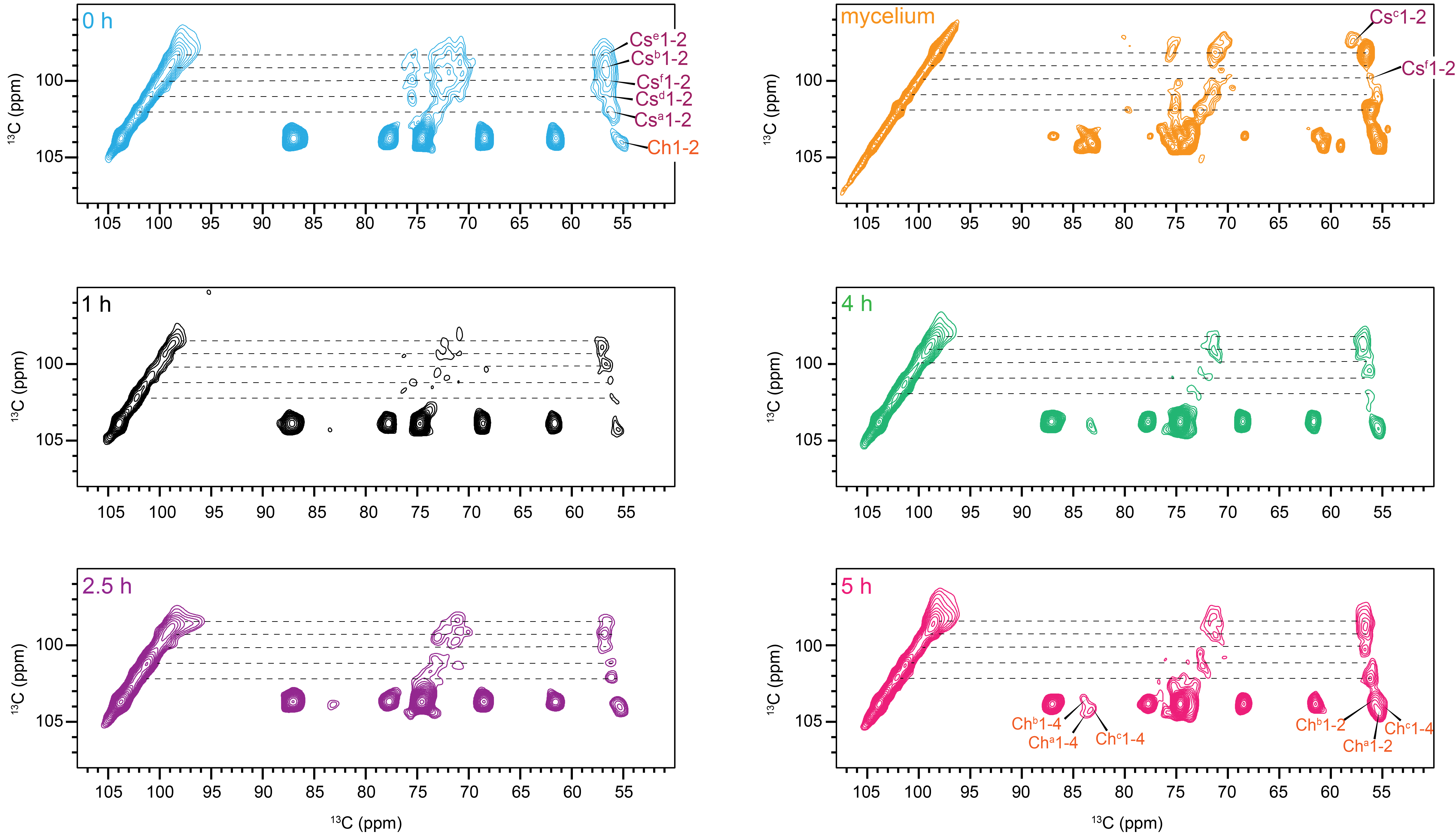

**Supplementary Figure 2.** **Polymorphic forms of chitosan in the germinating cell wall**. Zoomed view of C1 region from ^13^C-^13^C CORD spectra of *R. delemar* germinating conidia show structural transitions from dormant (0 h, blue) to germinations stages (1 h, black: 2.5 h, purple: 4 h, green: 5 h: pink) to fully developed mycelium (3 days, orange). All spectra from 1-5 h samples were obtained from cultures grown in labeled conidia and labeled media (LCLM). Multiple polymorphic forms are observed, with five distinct chitosan and three chitin forms detected in the isotropic conidia (pink), indicating dynamic remodeling during early germination. Chitosan type-c is exclusively detected in the 3-day culture, suggesting its emergence during mature hyphal development.

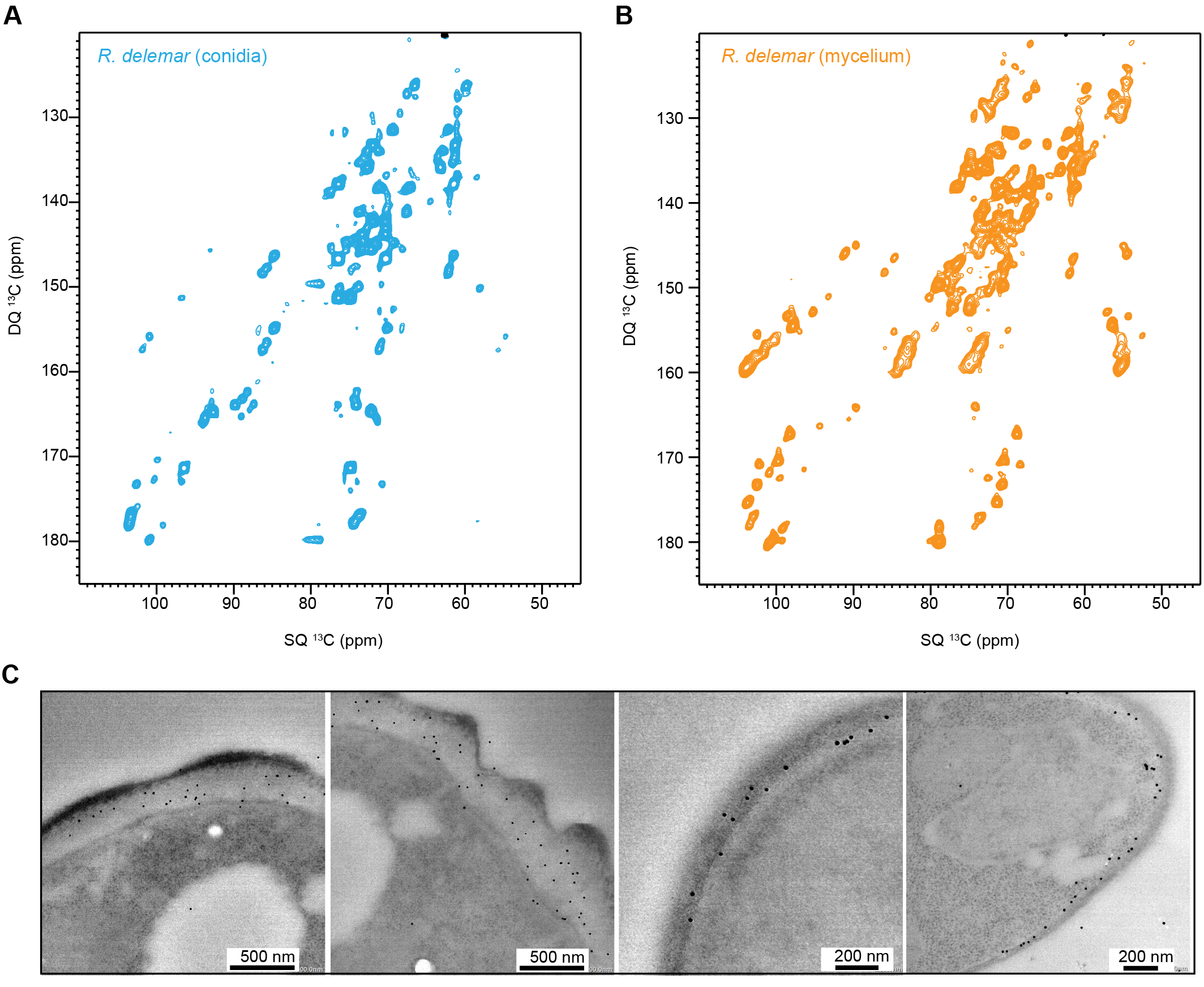

**Supplementary Figure 3.** **Mobile carbohydrate composition of *Rhizopus* conidia vs mycelium**. The full carbohydrate spectrum for (**A**) *Rhizopus* conidia is blue, and (**B**) the mycelium is orange. The spectra were collected at 800 MHz at 290 K with 15 kHz MAS. (**C**) The ConA-protein specificity shows positive binding, illustrating the localization of mannans.

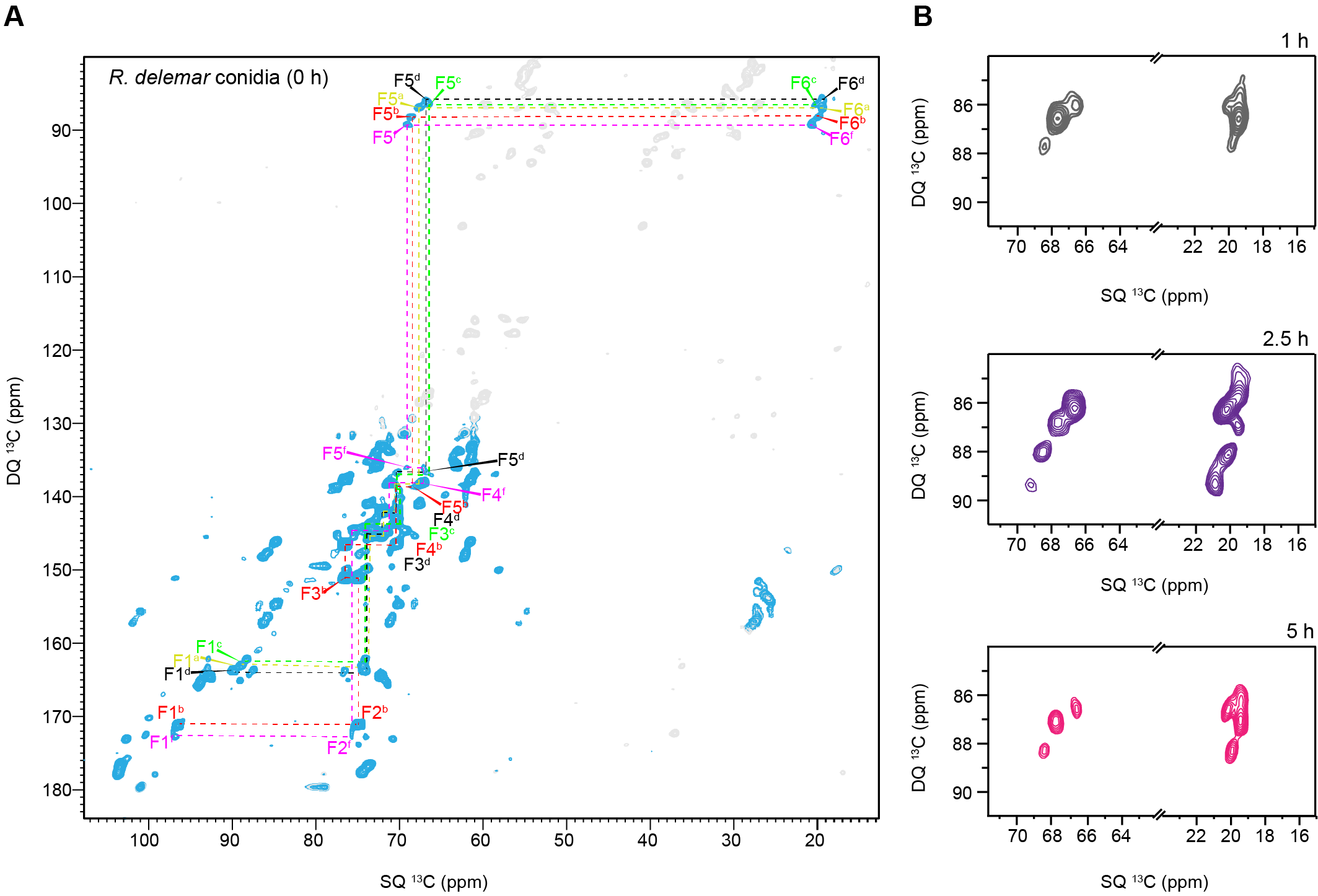

**Supplementary Figure 4.** **Polymorphism of fucose in *R. delemar* germinating conidia**. (**A**) Fucose and its polymorphic forms detected in the 2D DP-INADEQUATE spectrum of 0 h conidia. (**B**) Zoomed in view of F5-F6 carbon signals of Fucose units in R. delemar across different stages of germination (0 h, blue: 1 h, grey: 2.5 h, purple: 5 h, pink). Fucose type-f was not detected in 1 h and 5 h time points. The cultures were prepared using labeled conidia and labeled media.

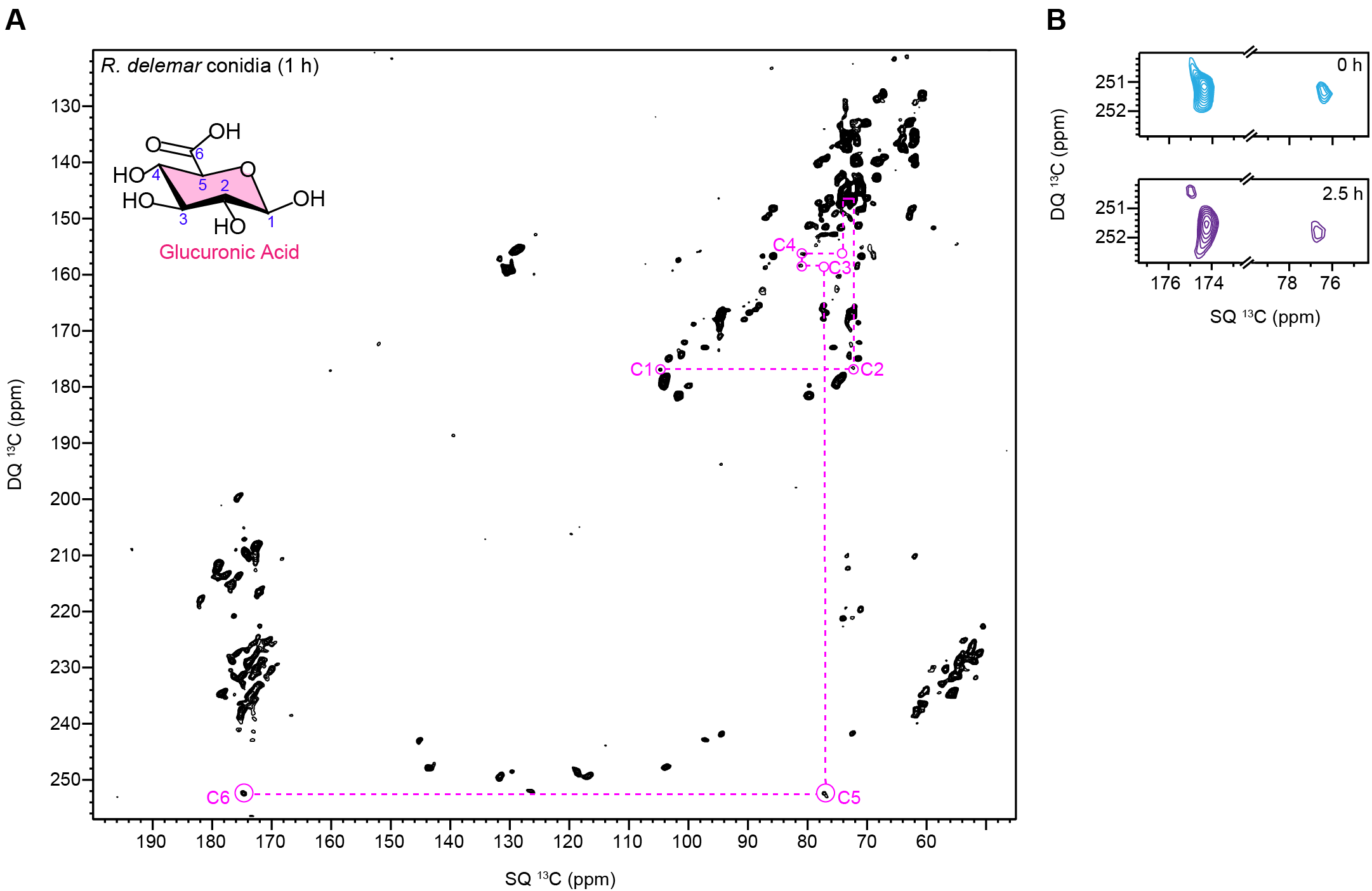

**Supplementary Figure 5.** **Glucuronic presence in *R. delemar* conidia**. (**A**) Glucuronic acid was identified in the 2D ^13^C DP INADEQUATE spectra of 1 h conidia, confirmed by the signature peaks at 175 ppm and 77 ppm. (**B**) Detection of GlcA units in dormant conidia (0h, blue) and swollen conidia (2.5 h, purple).

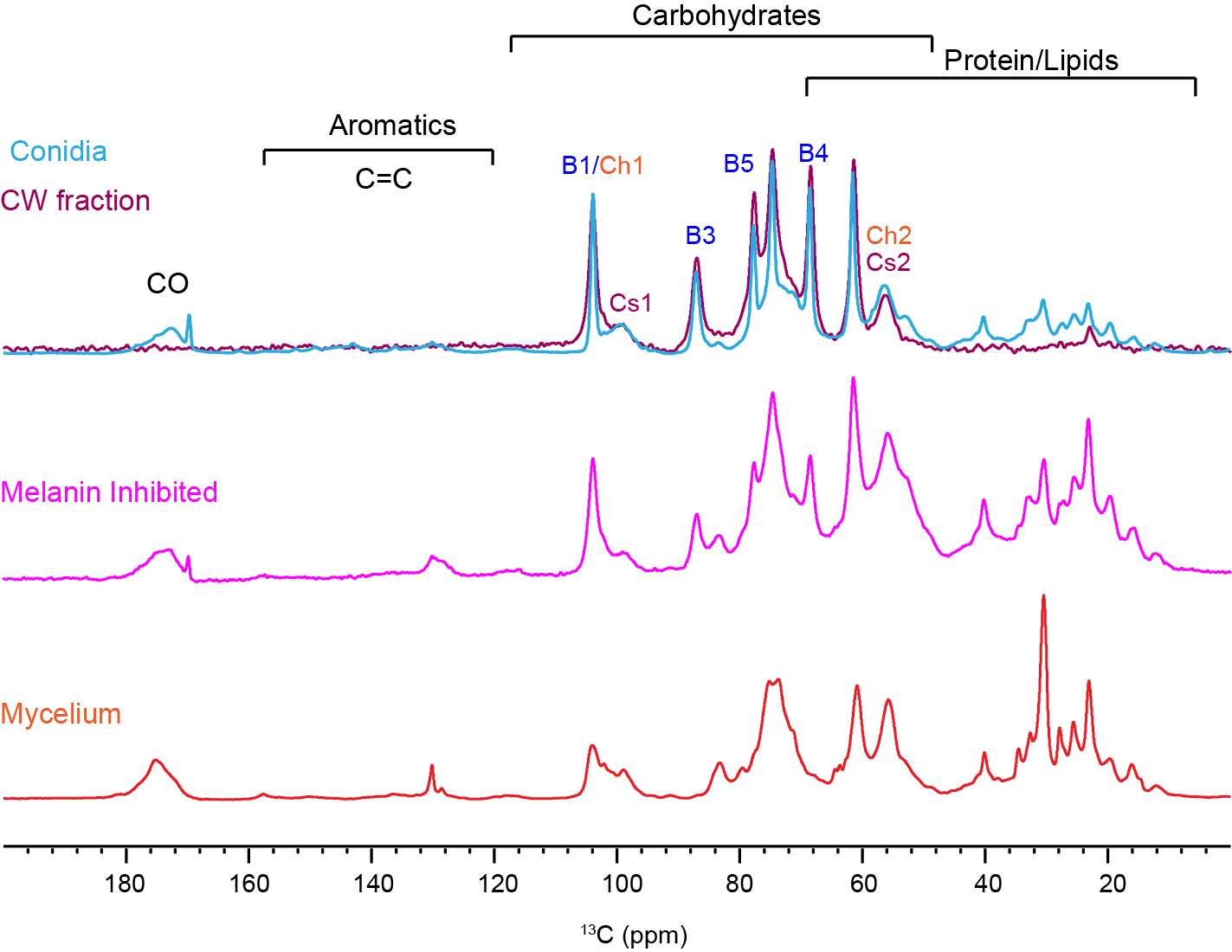

**Supplementary Figure 6. Comparison of cell walls under different conditions.** Overlay of intact conidia spectra (blue) with cell wall fraction (maroon). Spectra of melanin-inhibited cell walls, shown in magenta. Spectra of fully myceliated R. delemar cell walls. The spectra highlight resonances from carbonyl, aromatic, carbohydrate, and protein regions, revealing both conserved and condition-specific differences across developmental stages and melanin inhibition.

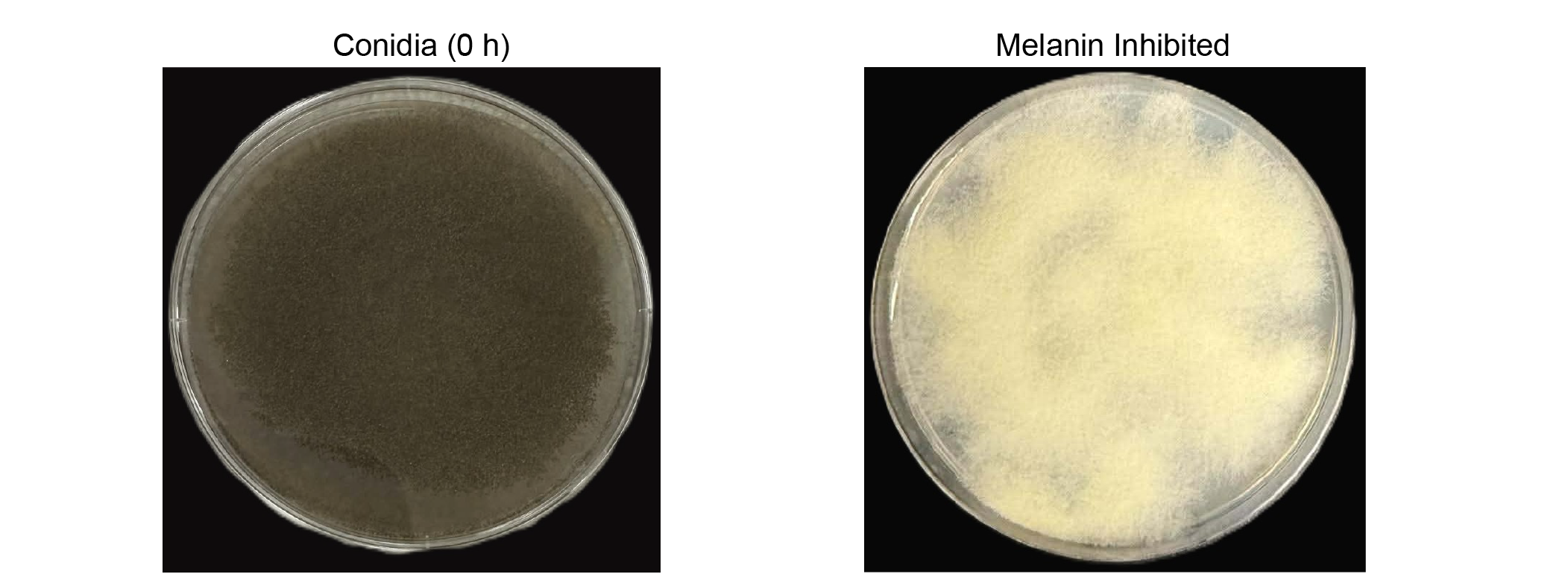

**Supplementary Figure 7. Conidia under control and melanin-inhibited conditions.** Untreated conidia displaying the typical dark pigmentation associated with melanin deposition (Left). Conidia grown in the presence of bathocuproinedisulfonic acid (BCS) show white colonies, indicating inhibition of melanin biosynthesis (Right).

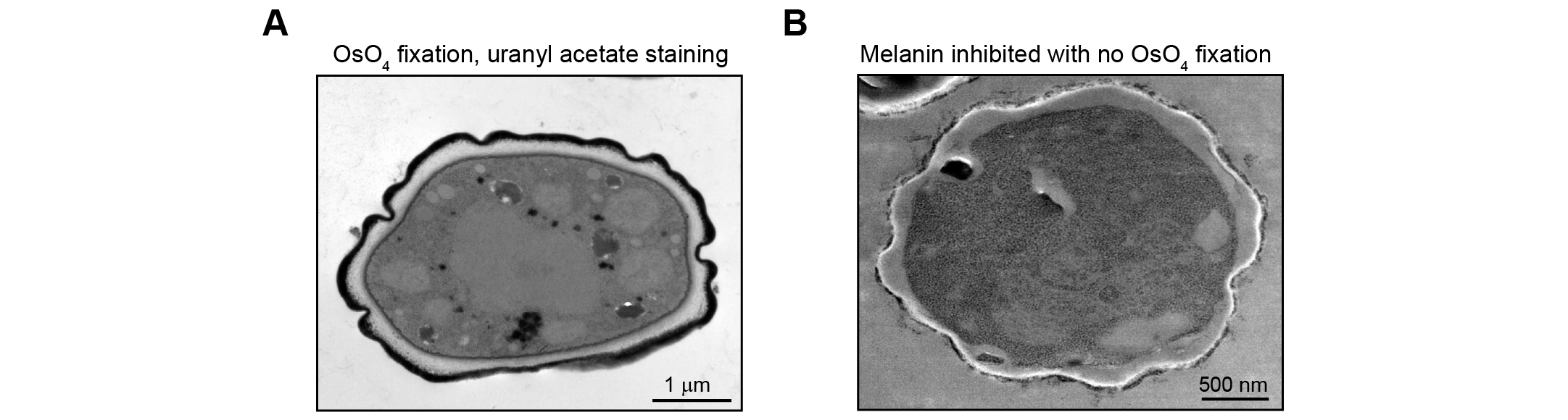

**Supplementary Figure 8. The cell wall ultrastructure in R. delemar conidia under different conditions.** (**A**) Dormant conidia display a multilayered cell wall architecture, consisting of an outer electron-dense layer followed by a granular layer and an inner electron-lucent layer. (**B**) Conidia treated with BCS to inhibit melanin biosynthesis, and prepared without osmium fixation, show loss of the electron-dense outer layer and the granular layer, with distinct electron-lucent layer. These structural changes indicate the contribution of melanin-rich components to outer wall architecture.

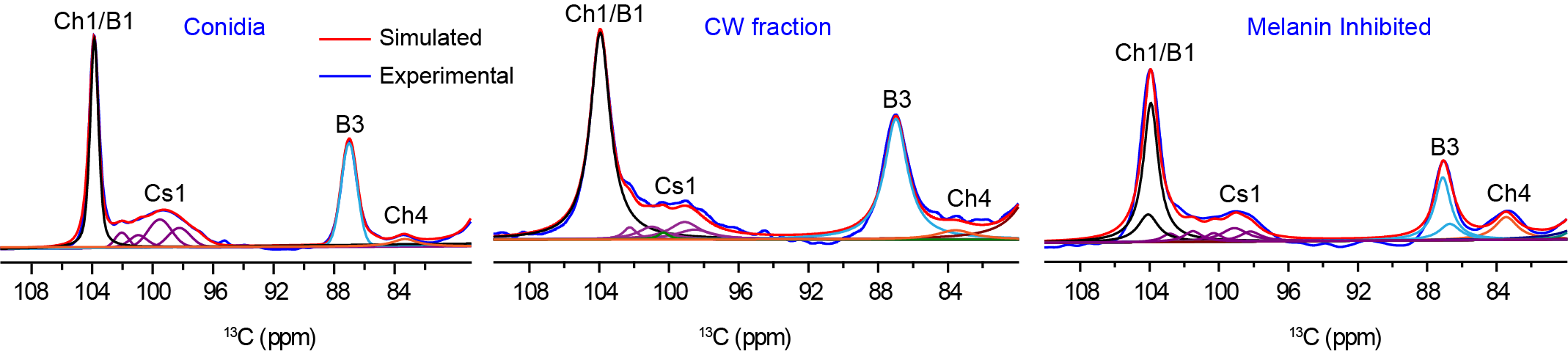

**Supplementary Figure 9. Deconvolution of polysaccharide of *R. delemer* under different conditions.** Deconvolution of region show contributions from chitin, β-1,3-glucan, and chitosan in conidia, cell wall fraction and BCS treatment (melanin inhibited). Cell wall fraction represents the higher contribution of β-1,3-glucan while melanin inhibition leads to increased chitin content, indicating significant restructuring of the cell wall scaffold.

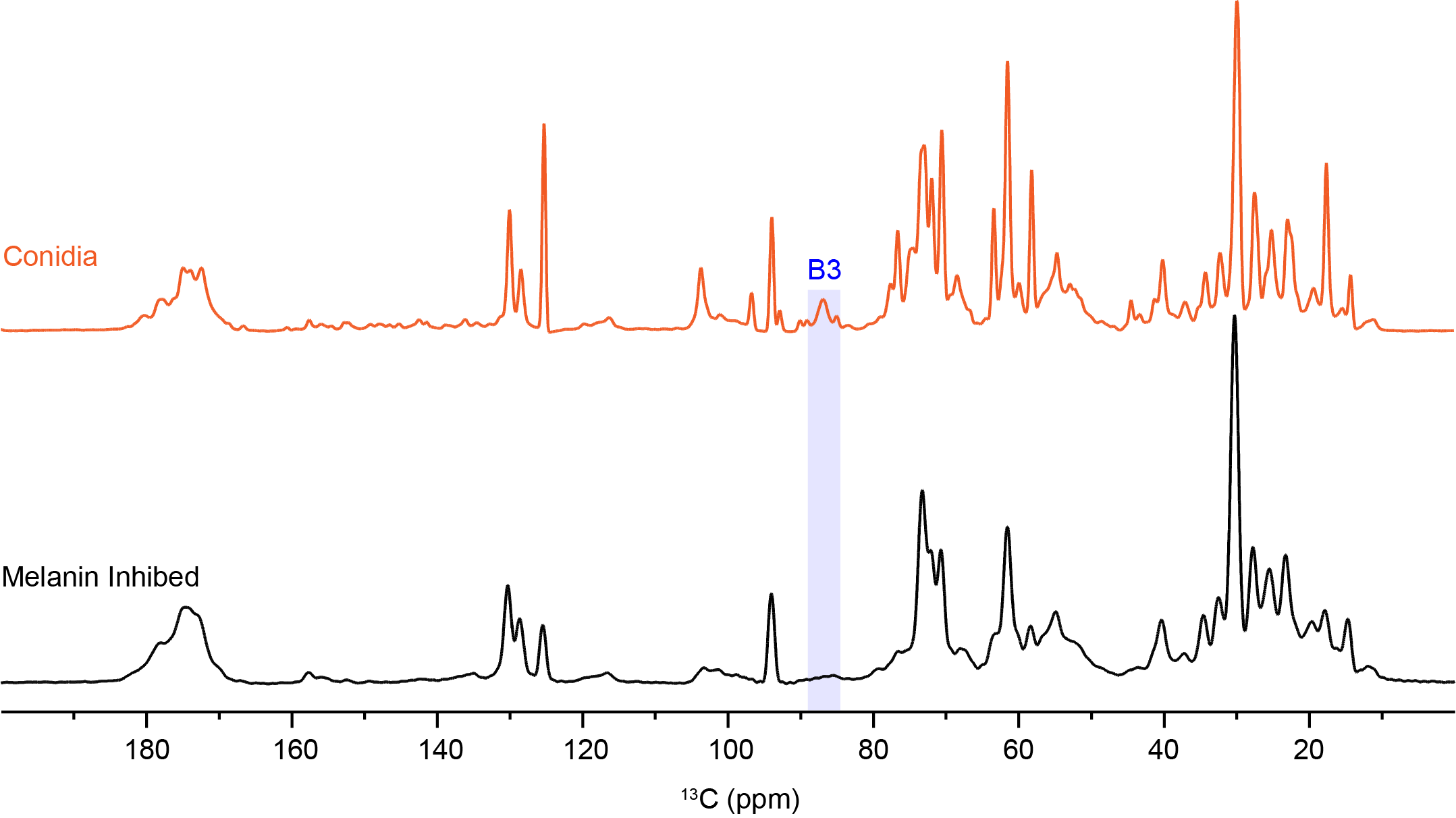

**Supplementary Figure 10:** **The effect of melanin inhibition on mobile β-1,3-glucan detection**. 2s DP ^13^C- spectra probing the mobile cell wall show that β-1,3-glucan signals are reduced under melanin-inhibited conditions (black spectra), in contrast to dormant conidia (orange spectra) where β-1,3-glucan remains detected by the glycosidic peak at B3 (highlighted by blue band).

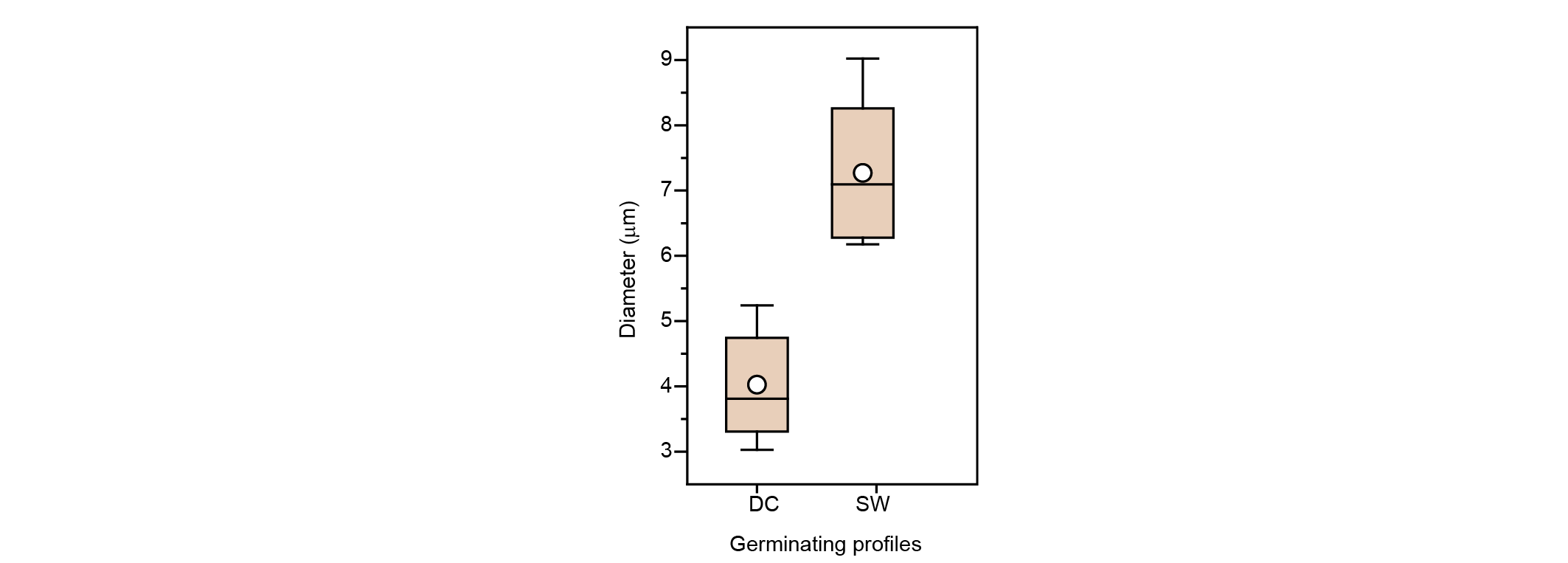

**Supplementary Figure 11**. **Box-and-whisker plot of conidial diameters.** Box-and-whisker plot showing the diameters of dormant and swollen conidia. The box represents the interquartile range (IQR), with whiskers extending to 1.5× IQR. The horizontal line indicates the median, and the open circle denotes the mean ± s.d. A total of n=9 cells were analyzed for each condition.

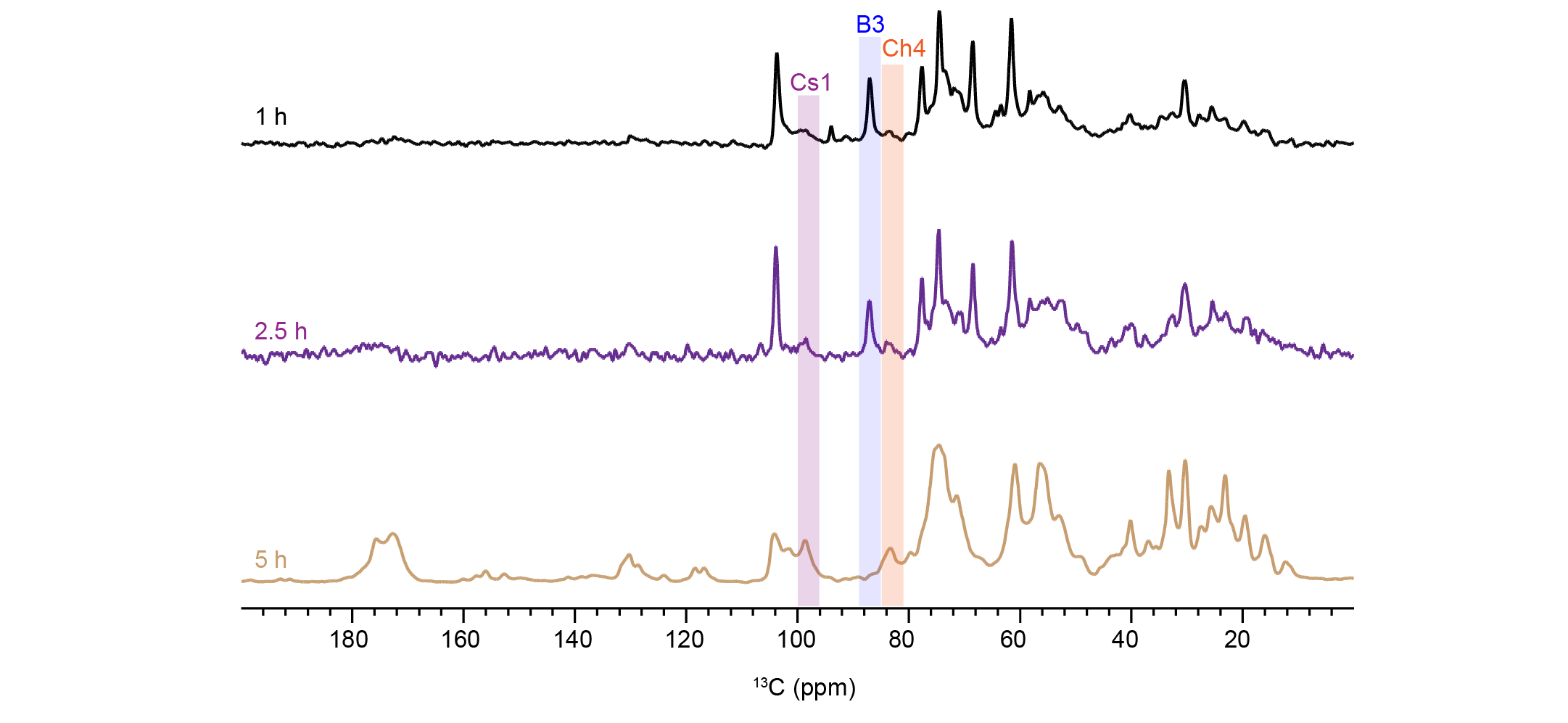

**Supplementary Figure 12. Neo-synthesized glucan composition during germination of *R. delemar*.** The germination stages span from early growth (1 h) to mycelial development (5 h). The 1D ^13^C CP spectra are color-coded as follows: 1 h (black, early stage/dormant-like conidia), 2.5 h (purple, swollen conidia), and 5 h (brown, germ tubes representing neo-synthesis). Progressive changes are observed across the spectra during germination. Signals corresponding to chitin and chitosan (83 ppm and ~99 ppm) increase over time, while the β-1,3-glucan resonance at ∼87 ppm decreases. Cultures were grown using unlabeled conidia in uniformly ^13^C-labeled media (unLCLM), enabling selective detection of neo-synthesized cell wall components.

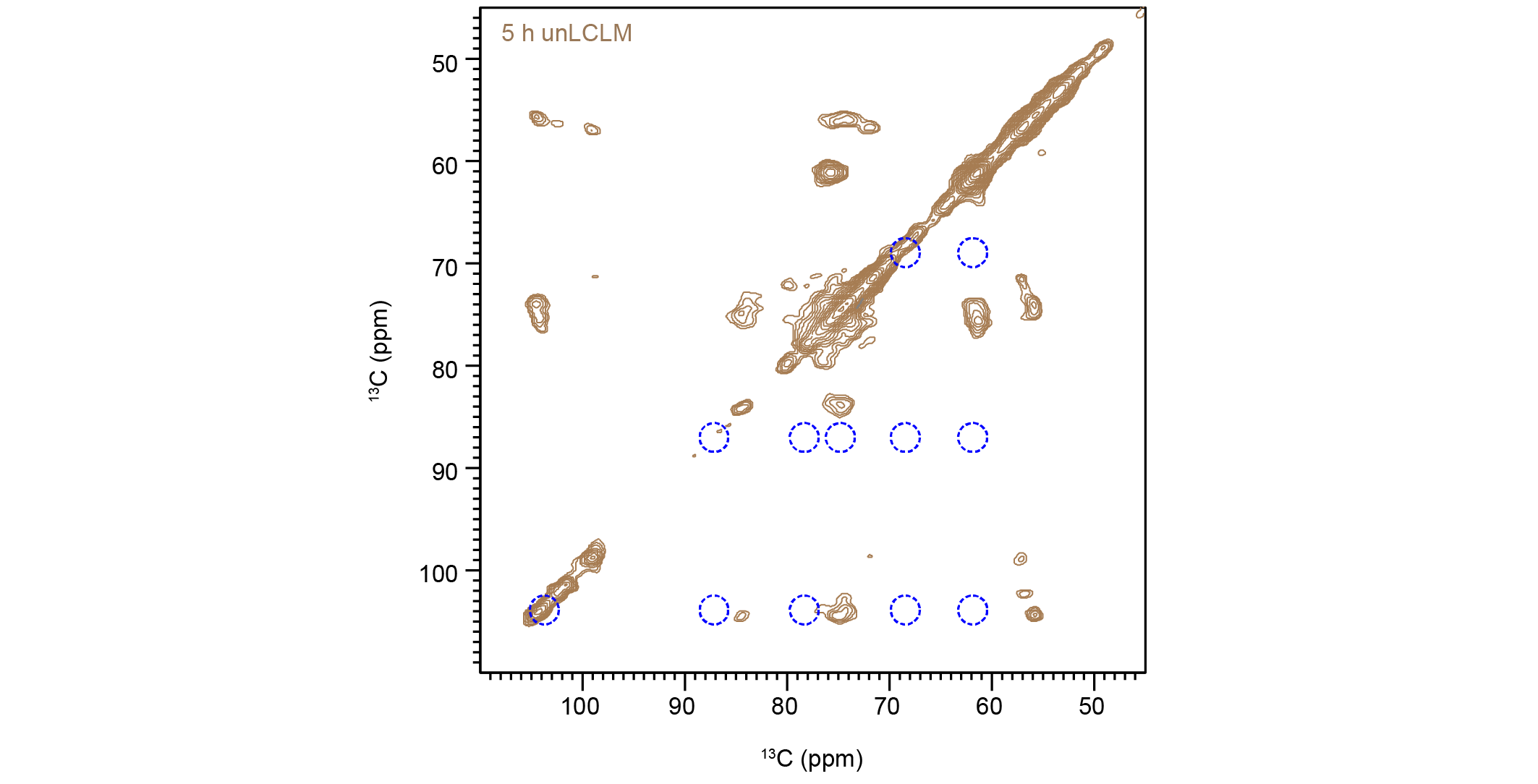

**Supplementary Figure 13. Rigid fraction of neo-synthesized molecules.** 2D ^13^C-^13^C CORD spectrum of 5 h germination, obtained using unlabeled conidia and ^13^C-labeled media (unLCLM) to selectively probe neosynthesized rigid molecules. The spectrum was obtained at 800 MHz at 290 K and 15 kHz MAS.

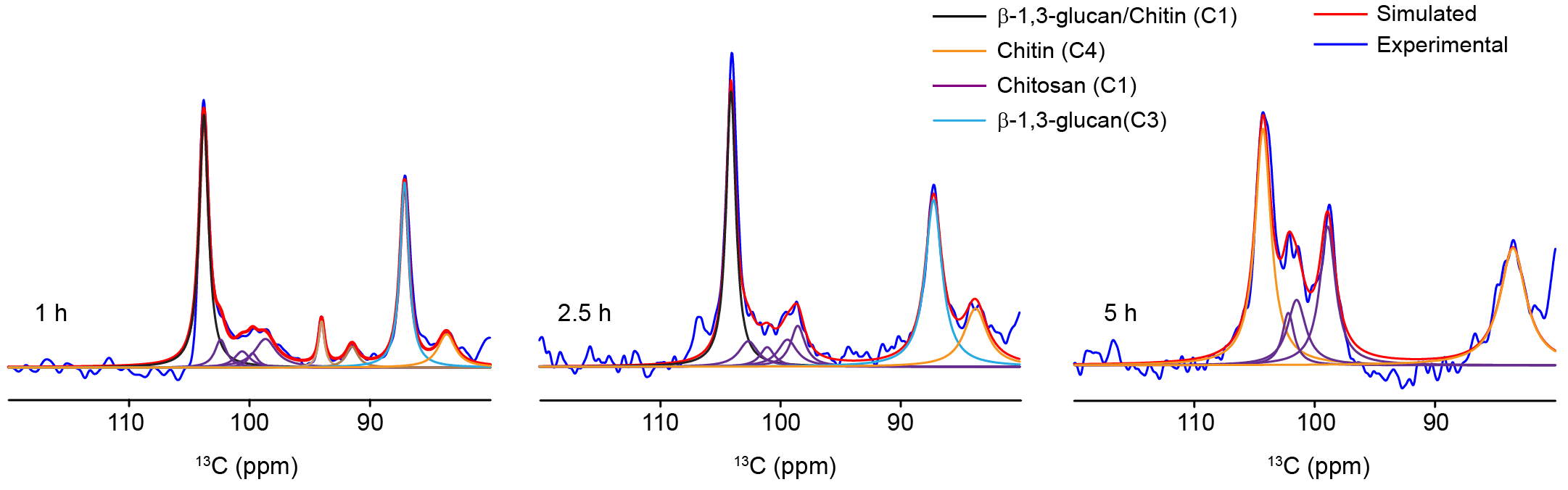

**Supplementary Figure 14. Deconvolution of neo-synthesized polysaccharides.** Deconvolution of 1D ^13^C CP spectra was performed on the C1 region for chitosan, the C1 and C3 regions for β-1,3-glucan, and the C1 and C4 regions for chitin. A progressive increase in chitin content is observed during germination (1 h to 5 h), accompanied by a decrease in β-1,3-glucan signals. Simulated spectra are shown in blue, and experimental spectra are shown in red. These results indicate dynamic remodeling of neo-synthesized polysaccharides during germination.

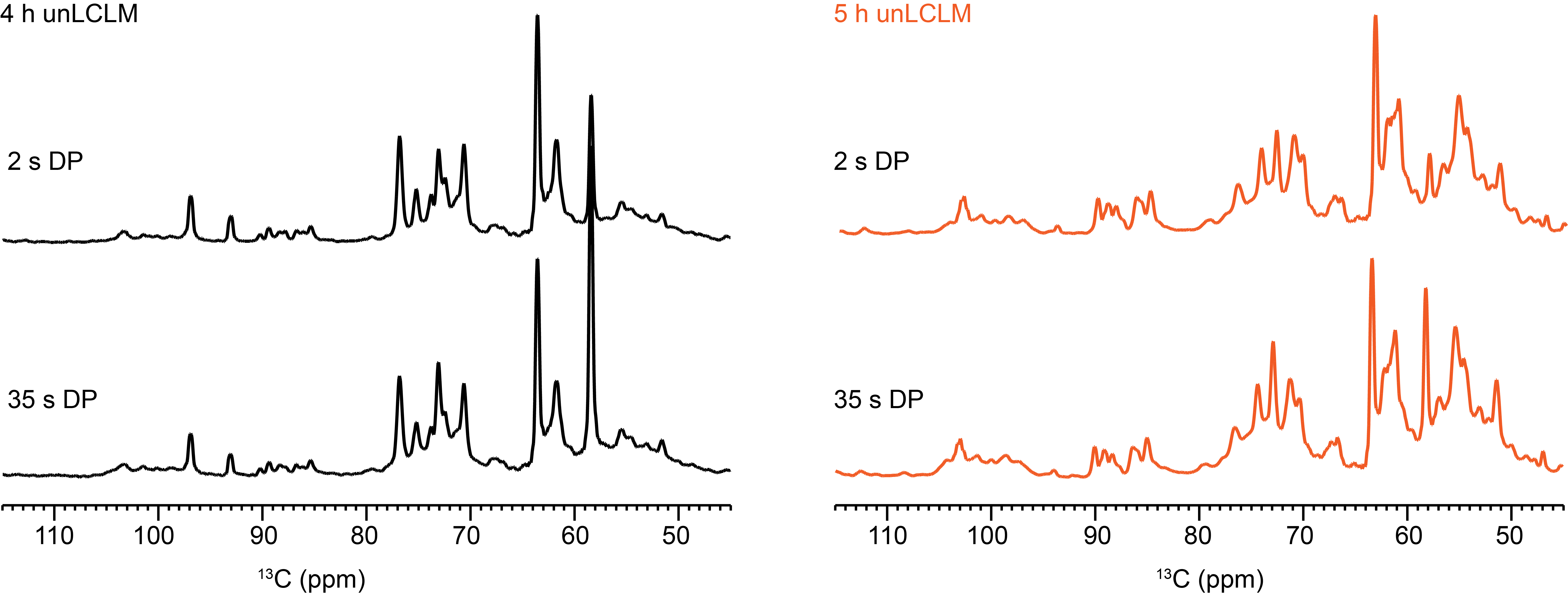

**Supplementary Figure 15.** **1D ^13^C spectra of neo-synthesized molecules without β-1,3-glucan**. ^13^C-DP spectra acquired with short recycle delay (2 s) selectively probing mobile region and long recycle delay (35 s) for quantitative detection of all molecules. Samples are color-coded by time point: black represents the 4-hour time point, and orange represents the 5-hour germination, grown from unlabeled conidia in ^13^C-labeled media (unLCLM).

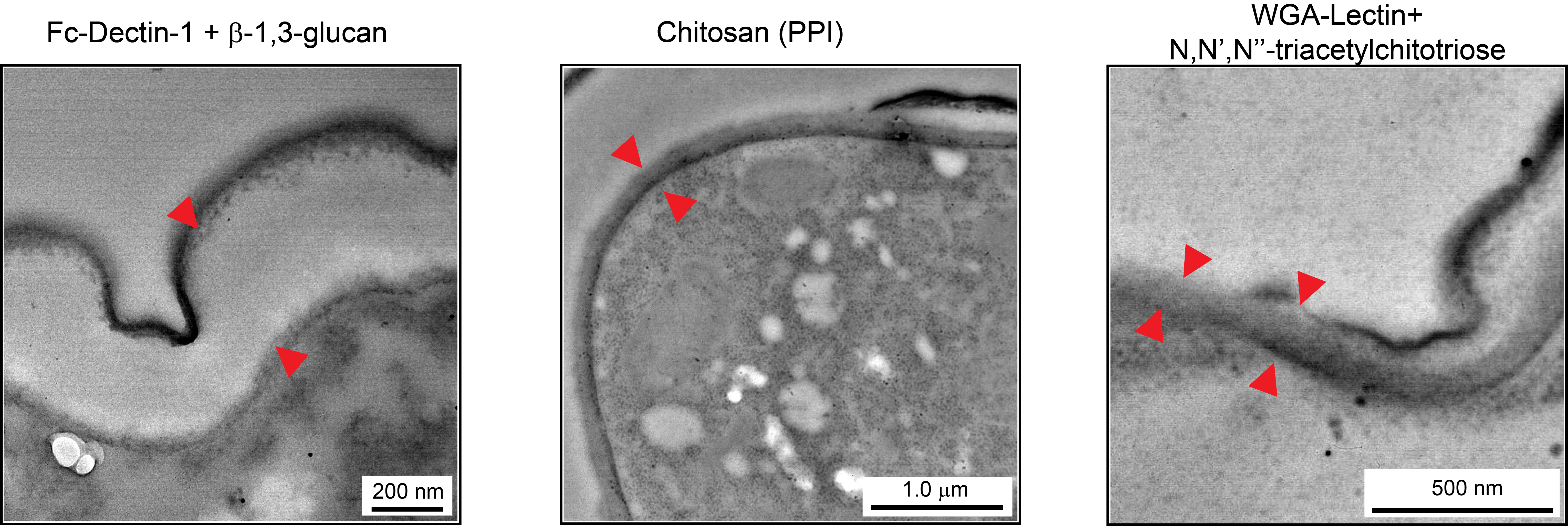

**Supplementary Figure 16. Control immunolabeling of β-1,3-glucan, chitosan, and chitin.** Control experiments were performed to validate the specificity of polysaccharide labeling**.** For β-1,3-glucan, purified glucan was incubated with Fc-Dectin-1 at a 1:1 ratio, followed by labeling with Protein A,10 nm gold (1:10 dilution). Chitosan controls were performed using pre-immune serum (1:100 dilution) and subsequently labeled with Protein A,10 nm gold (1:10), consistent with the chitosan-antiserum binding assay conditions. Chitin controls were prepared using a 1:500 dilution of chitin colloids and labeled with wheat germ agglutinin (WGA)-lectin gold conjugate (1:10 dilution). For specificity validation, WGA-lectin labeling was performed in the presence of 50 mM N,N',N'' Triacetylchitotriose as a competitive inhibitor, confirming specific binding to chitin. Red arrow shows absence of gold particles (black dots) binding.

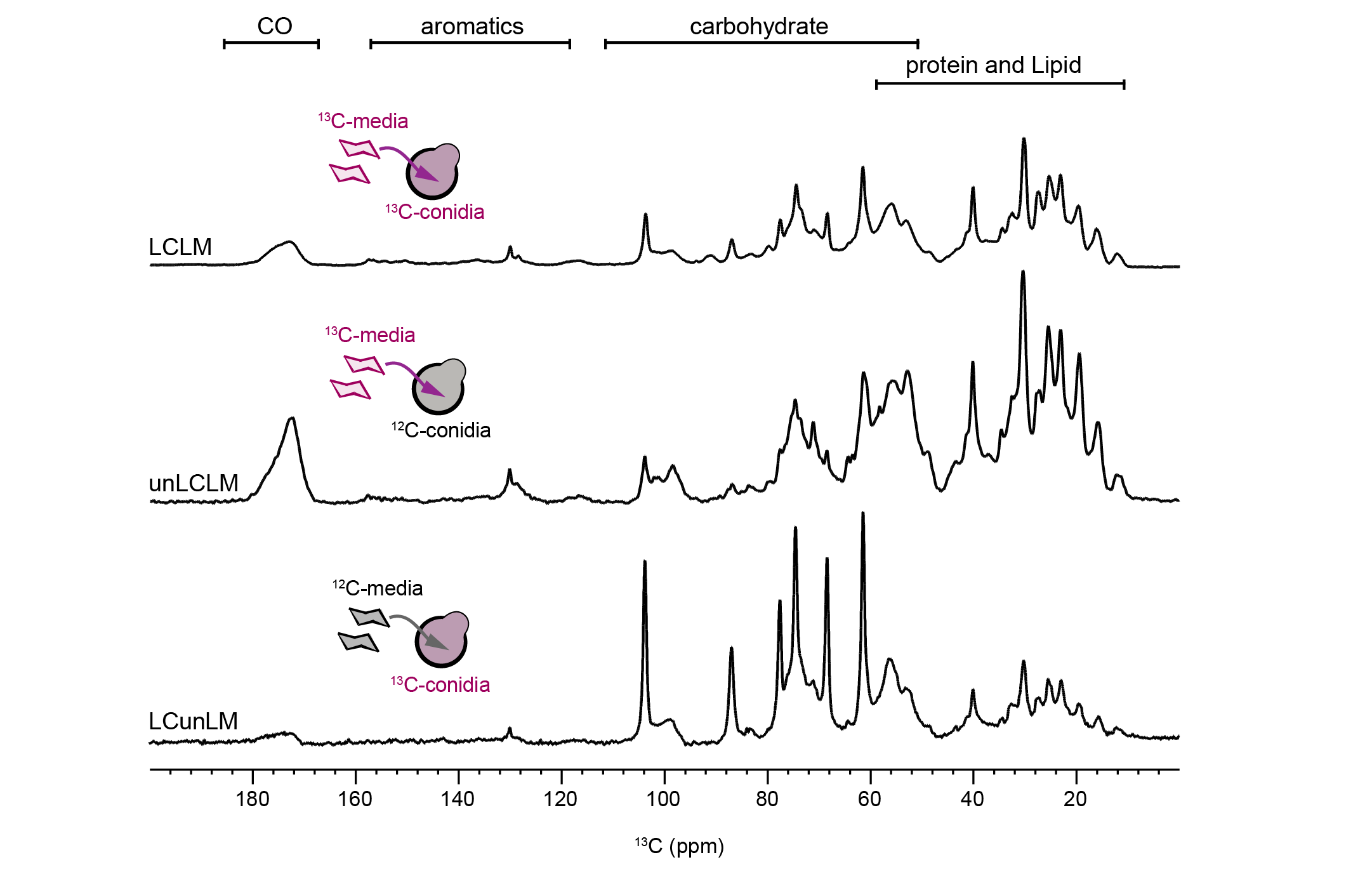

**Supplementary Figure 17. Cell wall labeling strategy and spectral comparison.** The labeling conditions include labeled conidia in labeled media (LCLM, top spectra), unlabeled conidia in labeled media (unLCLM, middle spectra), and labeled conidia in unlabeled media (LCunLM, bottom spectra), all acquired at the 4 h time point. The labeling scheme is illustrated in the model. Spectral regions corresponding to carbonyl (CO), aromatic, carbohydrate, and protein/lipid signals are indicated. The pre-existing cell wall fraction (LCunLM) shows sharper and more resolved carbohydrate resonances, consistent with a rigid β-glucan-chitin scaffold. In contrast, the neo-synthesized fraction (unLCLM) exhibits broader, more distributed signals, indicative of increased molecular mobility and enrichment in chitosan. The LCLM spectrum reflects the total cell wall pool, comprising both pre-existing and newly synthesized components.

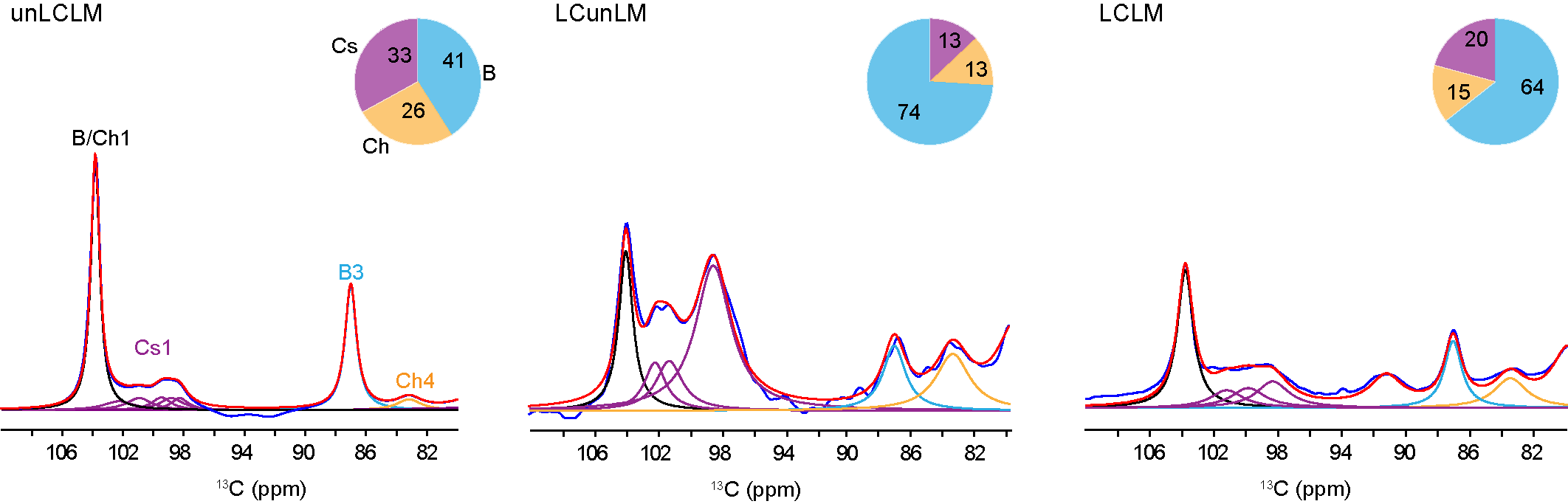

**Supplementary Figure 18. Polysaccharide composition at 4 h under different conditions.** Relative molar fractions of β-1,3-glucan (B), chitin (Ch), and chitosan (Cs) were determined from spectral analysis under three labeling conditions: newly synthesized fraction (unLCLM; unlabeled conidia in labeled media), pre-existing cell wall (LCunLM; labeled conidia in unlabeled media), and total pool (LCLM; labeled conidia in labeled media). Abbreviations denote carbon sites used for quantification (B3: β-1,3-glucan C3; Ch4: chitin C4; Cs1: chitosan C1; B/Ch: overlapping β-1,3-glucan and chitin C1). Pie charts show compositional distributions, revealing predominance of chitosan and chitin in the newly synthesized fraction with reduced β-1,3-glucan, enrichment of β-1,3-glucan and chitin in the pre-existing wall, and an intermediate profile in the total pool reflecting this transition. These results highlight selective remodeling of the fungal cell wall during growth.

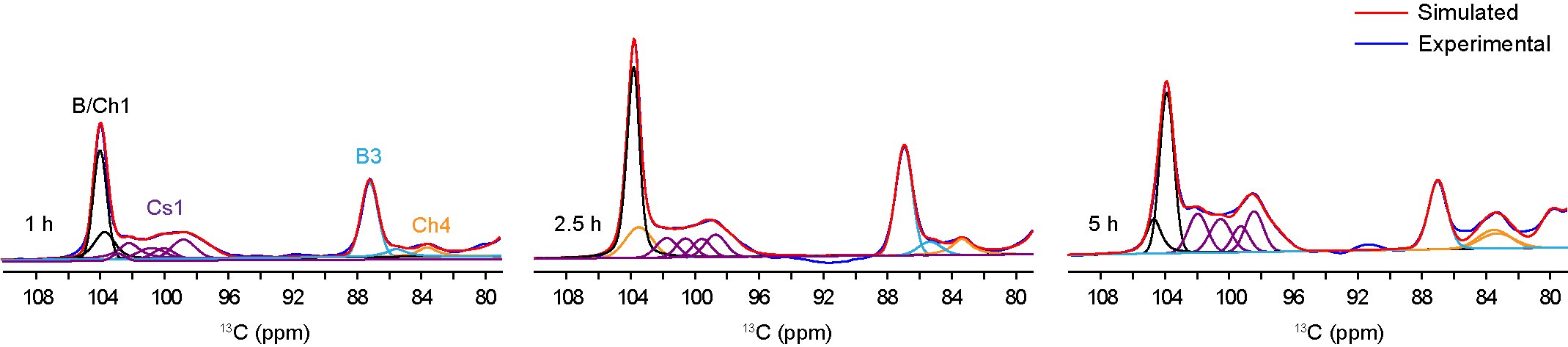

**Supplementary Figure 19. Germination stage-resolved polysaccharide composition.** Deconvolution of 1D ^13^C-CP spectra reveals stage-dependent changes in carbohydrate composition during germination in labeled conidia and labeled media (LCLM). Simulated spectra (red) closely match the experimental data (blue) and resolve contributions from individual polysaccharides. Colors and abbreviations denote polysaccharide types and the carbon sites used for relative quantification: B3 (cyan), C3 of β-1,3-glucan; Ch4 (orange), C4 of chitin; Cs1 (purple), C1 of chitosan; and B/Ch1, the overlapping C1 signal of β-1,3-glucan and chitin.

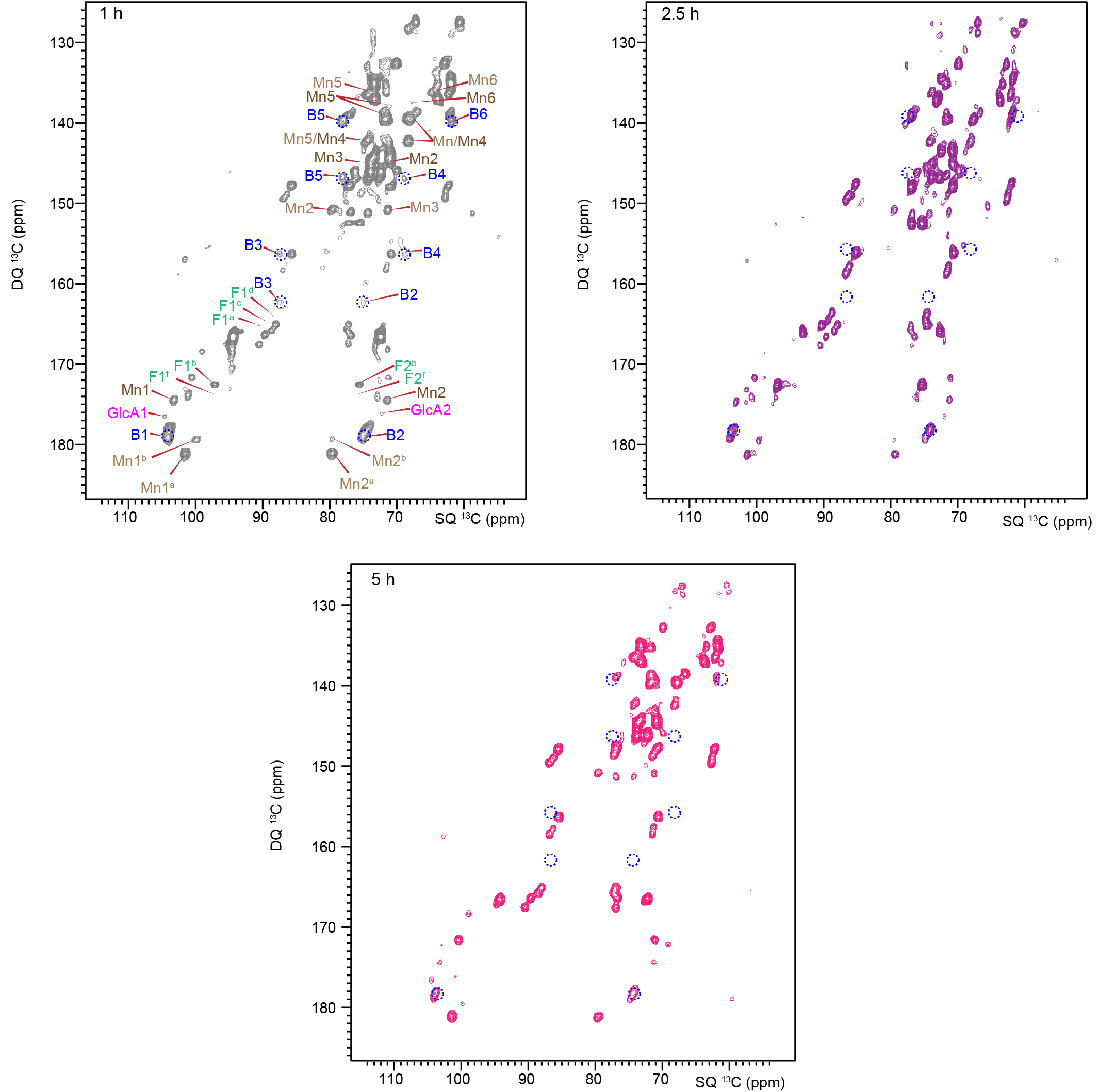

**Supplementary Figure 20. Mobile molecules of germinating conidia at different stages.** Spectra are shown for dormant conidia (grey, 1 h), swollen conidia (purple, 2.5 h), and germ tubes (pink, 5 h) for LCLM culture conditions. The spectra reveal progressive alterations in β-1,3-glucan spin-pairs (dotted blue circles), reflecting dynamic remodeling of mobile polysaccharide components during growth polarization.

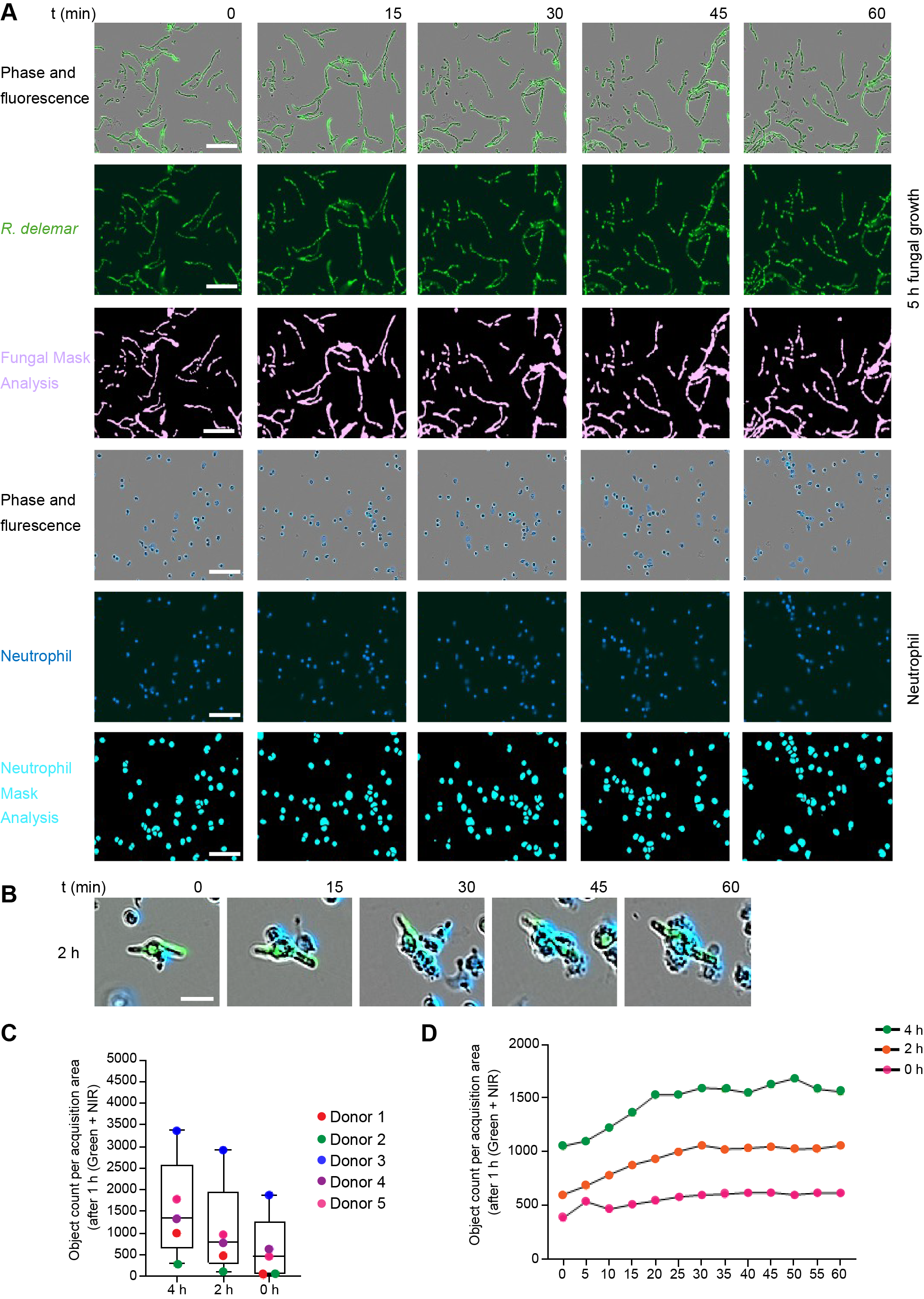

**Supplementary Figure 21.** **Neutrophil and *R. delemar* identification.** (**A**) Images showing staining and identification of *R. delemar* and neutrophils. Neutrophils were labeled with a near-infrared (NIR) tracker, and 5-hours growth *R. delemar* was stained with Sytotox Green. Live-cell imaging was performed for 60 min, with images acquired every 15 min. Image segmentation (mask analysis) identified *R. delemar* (pink) and neutrophils (blue). Scale bar, 100 µm. (**B**) Zoom in of picture highlighting the neutrophils binding kinetic on *R. delemar* at 2 h growing stage during 1 h. Scale bar, 20 µm. (**C**) Number of double-positive (NIR⁺/green⁺) objects per imaging field after 1 h of co-culture between human neutrophils (from five independent donors) and *R. delemar* resting conidia (0h) or fungi pre-grown in RPMI medium 2 and 4 h prior to interaction. Data is presented as mean ± s.e.m. from n = 5 independent donors. (**D**) Kinetics of neutrophil–fungus interactions, shown as total double-positive (NIR⁺/green⁺) objects per imaging field measured every 5 min over 1 h; 0 h (pink), 2 h (orange), 4 h (green).

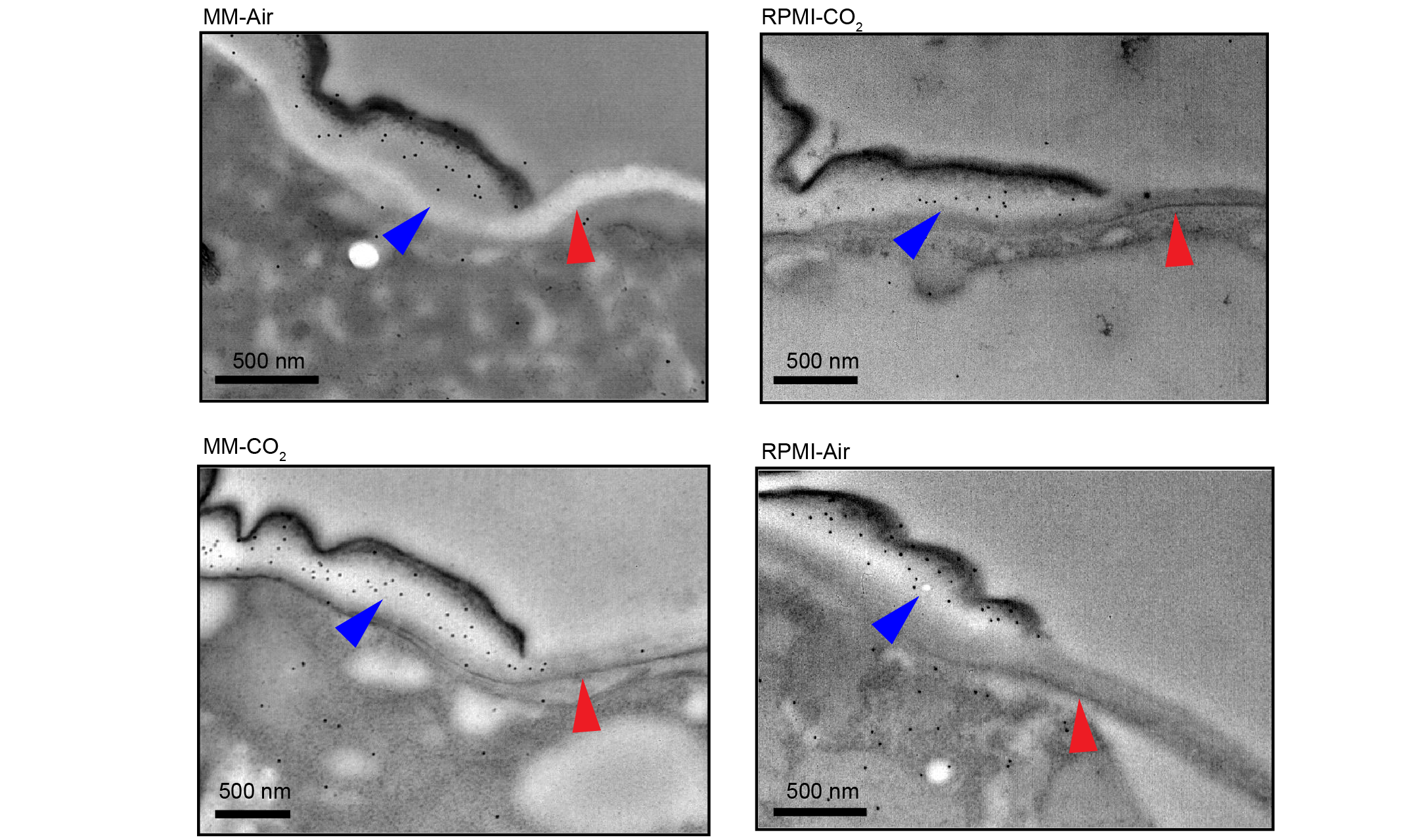

**Supplementary Figure 22. Absence of β-1,3-glucan in germ tubes under different growth conditions.** Fc–Dectin-1 binding was assessed in germinating conidia grown in RPMI under CO_2_-enriched (RPMI-CO_2_) and air (RPMI-Air) conditions, as well as minimal medium under CO_2_-enriched (MM-CO_2_) and air (MM-Air) conditions. Positive binding is observed in conidia (blue arrows), whereas no binding is detected in the germinating region or germ tube (red arrows). Scale bars represent 500 nm.

**Supplementary** **Table 1. Measurements of cell wall thickness and diameter of *R. delemar*.** All measurements are performed by ImageJ with n=120. Error bar: s.d. The average of the cell wall measurements with their standard deviation are listed below. The average diameter of the 9 cells with their standard deviation listed below.

| Cell wall thickness (nm) | |
| --- | --- |
| DC | 316±51 |
| SW | 229±62 |
| GT | 77±28 |
| Cell diameter (µm) | |
| DC | 4.0±0.7 |
| SW | 7±1 |

**Supplementary Table 2.** **Spectral deconvolution of 1D ^13^C CP spectra of *R. delemar* cell walls.** Deconvolution (79-110 ppm) was performed using DMfit for cell wall (CW) fraction isolated by boiling SDS/DTT and BCS-treated (melanin-inhibited) samples. C1 and C4 were used for chitin, C1 and C3 for β-1,3-glucan, and C1 for chitosan to estimate relative compositions.

| Polysaccharides | | ppm | Line width | | T_2_ (s) | Amplitude | Integral | Total % | |
| --- | --- | --- | --- | --- | --- | --- | --- | --- | --- |
| Conidia | | | | | | | | | |
| B1/Ch1 | 103.9 | | | 0.66 | 2.39 | 1.93E+07 | 4.22E+09 | |  |
|  | 104.1 | | | 0.66 | 1.59 | 1.93E+07 | 4.22E+09 | |  |
| B3 | 87.02 | | | 1.04 | 1.22 | 1.02E+07 | 3.32E+09 | | 54 |
| Ch4 | 83.42 | | | 2.09 | 0.44 | 872940 | 5.52E+08 | | 9 |
| Cs1 | 102 | | | 0.76 | 2.08 | 502680 | 1.10E+08 | | 37 |
|  | 100.97 | | | 0.86 | 1.84 | 620370 | 3.95E+08 | |  |
|  | 99.66 | | | 3.92 | 1.59 | 1846000 | 1.18E+09 | |  |
|  | 98.4 | | | 1.9 | 0.83 | 1180700 | 6.53E+08 | |  |
| CW Fraction | | | | | | | | | |
| B1/Ch1 | 103.94 | | | 1.38 | 1.14 | 942733.6 | 3.94E+08 | |  |
| Ch4 | 83.56 | | | 2.5 | 0.63 | 42230.03 | 2.93E+07 | | 7 |
| B3 | 86.98 | | | 1.6 | 0.99 | 549932.3 | 2.64E+08 | | 29 |
| Cs1 | 99.09 | | | 2.15 | 0.74 | 79639.67 | 5.14E+07 | | 64 |
|  | 98.52 | | | 2.48 | 0.64 | 44902.14 | 3.33E+07 | |  |
|  | 100.31 | | | 0.76 | 2.07 | 31541.56 | 7476253 | |  |
|  | 102.24 | | | 0.91 | 1.74 | 55590.61 | 1.55E+07 | |  |
|  | 100.98 | | | 1.86 | 0.85 | 58262.73 | 3.27E+07 | |  |
| Melanin Inhibited | | | | | | | | | |
| Ch1/B1 | 103.92 | | | 1.04 | 1.52 | 2379440 | 7.58E+08 | |  |
|  | 104.08 | | | 1.41 | 1.12 | 472382.7 | 2.01E+08 | |  |
| B3 | 87.1 | | | 1.11 | 1.43 | 1077265 | 3.64E+08 | |  |
|  | 86.7 | | | 1.46 | 1.08 | 278914.9 | 1.23E+08 | | 45 |
| Ch4 | 83.45 | | | 1.6 | 0.99 | 389478.3 | 1.80E+08 | | 28 |
| Cs1 | 100.3 | | | 1.11 | 1.43 | 147335.4 | 5.01E+07 | | 27 |
|  | 98.18 | | | 1.47 | 1.07 | 170205.7 | 7.67E+07 | |  |
|  | 99.01 | | | 1.23 | 1.29 | 103650.5 | 3.91E+07 | |  |
|  | 101.5 | | | 1.23 | 1.29 | 182474.8 | 6.86E+07 | |  |
|  | 99.1 | | | 1.84 | 0.86 | 230343.2 | 1.28E+08 | |  |

**Supplementary** **Table 3. ^13^C chemical shifts of rigid and mobile polysaccharides**. Superscripts are used to denote different allomorphs. Not applicable (/). Unidentified (-).

| Rigid Carbohydrate | | | C1 | C2 | C3 | | C4 | C5 | | | C6 | | CO | CH_3_ | | References |
| --- | --- | --- | --- | --- | --- | --- | --- | --- | --- | --- | --- | --- | --- | --- | --- | --- |
| Chitin | a | | 103.8 | 55.4 | 73.4 | | 83.5 | 75.4 | | | 60.5 | | 175.7 | 22.7 | | Kang et al., 2018^1^ Fernando et al 2021^2^ Cheng et.al 2024^3^  Chakraborty et al. 2021^4^ |
|  | b | | 103.6 | 55.6 | 73.2 | | 83.9 | 75.2 | | | 61.3 | | 174.5 | 22.8 | |  |
|  | c | | 104.4 | 55.2 | 73.4 | | 83.7 | 75.7 | | | 60.8 | | 176.7 | 23.2 | |  |
| Chitosan | a | | 102.1 | 56.0 | 72.6 | | 79.5 | 75.3 | | | - | | / | / | |  |
|  | b | | 98.6 | 56.6 | 71.6 | | 76.6 |  | | | - | | / | / | |  |
|  | e | | 101.0 | 55.9 | 72.4 | | 76 | 73.9 | | | - | | / | / | |  |
|  | f | | 97.3 | 56.7 | 71.3 | | - | - | | | - | | / | / | |  |
|  | g | | 100.2 | 56.7 | 72.3 | | - | - | | | - | | / | / | |  |
| β-1,3-glucan |  | | 103. 8 | 74..6 | 86.9 | | 68.5 | 77.7 | | | 61.5 | | / | / | |  |
| Mobile Carbohydrate | | Types | | C1 | | C2 | | | C3 | C4 | | C5 | | | C6 | References |
| α-Mn^1,2^ | | a | | 101.4 | | 79.4 | | | 71.5 | 67.8 | | 74.2 | | | 62.5 | Dickwella et al. 2024^5^ |
|  | | b | | 99.9 | | 79.5 | | | - | - | | - | | | - |  |
| α-Mn^1,6^ | | - | | 103.2 | | 71.3 | | | 73.7 | 67.9 | | 71.3 | | | 67.3 |  |
| β-1,3-glucan (B) | | - | | 104.1 | | 74.3 | | | 85.6- | 70.6 | | 77.2 | | | 62.0 |  |
| Fucose (F) | | a | | 90.8 | | 74.2 | | | 72.4 | 71.3 | | 68.6 | | | 19.8 |  |
|  |  | b | | 97.5 | | 74.8 | | | 77.8 | 69.5 | | 68.1 | | | 21.1 |  |
|  |  | c | | 90.7 | | 74.4 | | | 70.7 | 73.3 | | 66.1 | | | 20.7 | Cheng et. al 2024^3^ |
|  |  | d | | 94.5 | | 72.2 | | | 72.3 | 70.1 | | 67.4 | | | 19.9 |  |
|  |  | i | | 97.2 | | 75.5 | | | 77.1 | 70.7 | | 69.2 | | | 20.3 | Furevi, et. al 2022^6^ |
| Glucose | | a | | 93.2 | | 72.8 | | | 74.0 | 71.2 | | 70.9 | | | 63.8 |  |
|  |  | b | | 94.9 | | 72.3 | | | 73.8 | 71.1 | | 77.2 | | | 62.1 | Dickwella et al 2025^7^ |
| Glucuronic Acid  (GlcA) | | - | | 103.8 | | 73.4 | | | 75.2 | 80.9 | | 77.3 | | | 174.6 | Ruiter et.al 1992^8^ |

**Supplementary** **Table 4.** **Solid-state NMR experimental parameters for cell wall characterization**. T = sample temperature 290 K for all experimental conditions; B_0_ = magnetic field of 18.8T; ν_MAS_ = MAS frequency; ns = number of scans; d_1_ = recycle delay between scans; t_1, max_ = maximum t_1_ evolution time (for indirect dimension); t_1, inc_ = increment for t_1_ (for indirect dimension) evolution time; τ_dw_ = dwell time during direct FID acquisition; τ_acq_ = maximum acquisition time during direct FID detection; τ_XY_ = cross-polarization contact time during CP from channel X to channel Y; ν_1H, dec_ = dipolar decoupling field strength.

| Samples | Experiment | NS | d1  (s) | t_1, max_  (ms) | t_1, inc_  (µs) | τ_acq_  (ms) | τ_HC_  (ms) | τ_mix_  (ms) | ν_1H dec_ (kHz) |
| --- | --- | --- | --- | --- | --- | --- | --- | --- | --- |
| LCLM | 1D ^13^C CP | 1024 | 2 |  |  | 16/  18 | 1 |  | 83 or 72.33 |
|  | 1D ^13^C DP | 512 | 2 |  |  |  |  |  | 100 |
|  | 1D ^13^C DP | 512 | 35 |  |  |  |  |  |  |
|  | 2D CORD | 32 | 2 | 7.5 | 25 | 14/  18 | 1 | 53 τ_CORD_ | 83 or 71 |
|  | 2D DP J-INADEQUATE | 32 | 1.5 | 10 | 20 or 22 | 18/  19 |  |  | 83 |
| unLCLM | 1D ^13^C CP | 1K-16K | 1.8-2.0 |  |  |  |  |  |  |
|  | 1D ^13^C DP | 1024 | 2 |  |  |  |  |  |  |
|  | 1D ^13^C DP | 1024 | 35 |  |  |  |  |  |  |
|  | 2D CORD | 64 | 2 | 7.5 | 25 | 14 | 1 | 53 τ_CORD_ | 83 or 72.33 |

**Supplementary Table 5.** **Spectral deconvolution of 1D ^13^C-CP spectra of *R. delemar*.** Deconvolution was performed using DMfit software6 from 110-79 ppm. The relative abundances represent the kinetics of the carbohydrate with germination.

| Carbohydrates | ppm | Line width | T2 | Amplitude | Integer | % |
| --- | --- | --- | --- | --- | --- | --- |
| 1h | | | | | | |
| Ch1 | 103.65 | 1.53 | 1.03 | 1867180.89 | 7.38E+08 |  |
| B1/Ch1 | 103.92 | 0.98 | 1.61 | 7448091.74 | 1.57E+09 |  |
| B3 | 87.13 | 1.23 | 1.29 | 5096488.53 | 1.34E+09 | 45 |
|  | 86.47 | 1.53 | 1.03 | 535740.13 | 2.48E+08 |  |
| Ch4 | 83.54 | 1.41 | 1.12 | 633791.16 | 2.67E+08 | 8 |
| Cs1 | 102.11 | 1.6 | 0.99 | 1100168.96 | 5.31E+08 | 48 |
|  | 97.48 | 1.9 | 0.83 | 866721.14 | 3.53E+08 |  |
|  | 100.73 | 1.96 | 0.81 | 741658.23 | 3.12E+08 |  |
|  | 99.96 | 1.47 | 1.07 | 720216.98 | 2.27E+08 |  |
|  | 98.73 | 1.72 | 0.92 | 1300671.54 | 4.79E+08 |  |
| 2.5 h | | | | | | |
| Ch/B1 | 103.89 | 0.92 | 1.72 | 9848375.65 | 2.27E+09 |  |
|  | 103.51 | 1.97 | 0.8 | 1580027.56 | 6.66E+08 |  |
| B3 | 87.01 | 1.23 | 1.29 | 5591301.75 | 1.47E+09 | 55 |
|  | 85.9 | 1.69 | 0.94 | 694022.73 | 2.52E+08 |  |
| C4 | 83.39 | 1.41 | 1.12 | 829033.26 | 3.48E+08 | 11 |
| Cs1 | 100.61 | 1.35 | 1.17 | 981503.45 | 2.84E+08 | 34 |
|  | 99.59 | 1.35 | 1.17 | 946772.71 | 2.74E+08 |  |
|  | 101.77 | 1.53 | 1.03 | 1005538.83 | 3.31E+08 |  |
|  | 98.73 | 1.53 | 1.03 | 1160571.64 | 3.82E+08 |  |
| 5 h | | | | | | |
| B1/Ch1 | 103.89 | 1.04 | 1.52 | 9058115 | 2.03E+09 |  |
|  | 104.7 | 1.11 | 1.43 | 1941160 | 5.97E+08 |  |
| B3 | 87.01 | 1.35 | 1.17 | 3865024 | 1.18E+09 | 26 |
| Ch4 | 83.51 | 2.33 | 0.68 | 1048089 | 5.24E+08 | 22 |
|  | 83.33 | 2.52 | 0.63 | 843022.6 | 4.54E+08 |  |
| Cs1 | 102.82 | 0.98 | 1.61 | 1245646 | 2.62E+08 | 52 |
|  | 98.46 | 1.35 | 1.17 | 2299765 | 6.65E+08 |  |
|  | 101.96 | 1.41 | 1.12 | 2212781 | 6.69E+08 |  |
|  | 100.55 | 1.53 | 1.03 | 1889589 | 6.21E+08 |  |
|  | 99.29 | 1.23 | 1.29 | 1467234 | 3.86E+08 |  |

**Supplementary** **Table 6. Molar composition of rigid polysaccharides in cell walls**. Molar percentages of rigid cell-wall polysaccharides are estimated using integrals (volume) of cross peaks in 2D ^13^C-^13^C 53-ms CORD. The molar percentages of rigid cell-wall polysaccharides are estimated by using well-resolved peaks of 2D space through ^13^C-^13^C CORD. (-) represents not detected. The data for 3 d mycelium is adopted from Cheng.*et.al* 2024^3^. (m) represents minor components. The neo-synthesis composition was calculated by convoluting from 1D CP spectra.

| Carbohydrates | Types | Dormant Conidia | Mycelium (3 d) |
| --- | --- | --- | --- |
| β-1,3-glucan (B) | - | 61 ± 0.4 | 5 ± 0.6 |
| Chitin  (Ch) | a/b/c  d | 5 ± 0.1  - | 43 ± 2  1 ±0.2 |
| Chitosan  (Cs) | a  b  c  d  e  f | 6 ± 0.1  11± 0.1  -  5± 0.1  5 ± 0.1  7± 0.1 | 13 ± 3  16 ± 2  6 ± 1  10 ± 1  m  - |

| Carbohydrates | Amplitude | ppm | Line width | T2 | Integral | % |
| --- | --- | --- | --- | --- | --- | --- |
| 1h | | | | | | |
| B/Ch1 | 4991900 | 103.8 | 0.93 | 1.7 | 1.45E+09 | 65 |
| B3 | 3658800 | 87.14 | 0.9 | 1.76 | 1.03E+09 | 23 |
| Ch4 | 660620 | 83.65 | 1.75 | 0.91 | 3.59E+08 |  |
| Cs | 551400  319880  282180  565250 | 102.4  100.62  99.75  98.69 | 1.16  1.55  1  2.13 | 1.36  1.02  1.58  0.74 | 2.01E+08  1.55E+08  8.84E+07  3.74E+08 | 13 |
| 2.5 h | | | | | | |
| B/Ch1 | 522793.2 | 104.04 | 0.96 | 1.64 | 1.58E+08 | 60 |
| B3 | 316658.3 | 87.18 | 1.69 | 0.94 | 1.67E+08 |  |
| Ch4 | 110523.5 | 83.69 | 2.3 | 0.69 | 7.91E+07 | 28 |
| Cs1 | 48533.05  37081.11  51436.92  77619.61 | 102.6  101.03  99.35  98.49 | 1.93  1.45  1.93  1.45 | 0.82  1.09  0.82  1.09 | 2.91E+07  1.68E+07  3.09E+07  3.51E+07 | 12 |
| 5 h | | | | | | |
| Ch1 | 996950 | 104.3 | 1.6 | 0.99 | 4.99E+08 | 46 |
| Ch4 | 493960 | 83.52 | 2.6 | 0.61 | 3.99E+08 |  |
| Cs1 | 221010  275730  585150 | 102.2  101.51  98.93 | 1.2  1.84  1.58 | 1.32  0.86  1 | 8.32E+07  1.59E+08  2.89E+08 | 54 |

Neo-synthesis

**Supplementary** **Table 7. Molar composition of mobile polysaccharides in cell walls**. The molar percentages of mobile cell-wall polysaccharides are estimated from well-resolved in 2D J-based ^13^C-^13^C DP-INADEQUATE spectra. Quantification was performed using C1 and C2 correlations for most polysaccharides, except for fucose, where C5 and 6 resonances were used and Glucuronic acid (GlcA ) was identified was CO-C5 resonances. (-) represented the non-detected signals.

| Carbohydrates | Dormant conidia | Swollen conidia | Germinating tube | Mycelium |
| --- | --- | --- | --- | --- |
| B | 25 | 28 | 20 | 4 |
| Mn^1,2 a^ | 15 | 13 | 23 | 16 |
| Mn^1,2b^ | 3 | 4 | 2 | 3 |
| Mn^1,2c^ | - | - | - | 2 |
| Mn^1,6^ | 6 | 5 | 4 | 7 |
| Gal(a-e) | 4 | 7 | 19 | 17 |
| F^i^ | 7 | 5 | 0 | 0 |
| F^a^ | 5 | 6 | 10 | 3 |
| F^b^ | 9 | 7 | 6 | 1 |
| F^d^ | 8 | 7 | 6 | 1 |
| F^c^ | 6 | 9 | 10 | 1 |
| F^e^ | 0 | 0 | -- | 2 |
| Cs^a,b^ | 3 | 1 | - | 8 |
| Cs^e^ | 4 | 3 | - | - |
| Ch^d^ | - | - | - | 15 |
| Ch^a,b^ | - | - | - | 6 |
| GlcA | 5 | 5 | - | - |
| GalNAc | - | - | - | 10 |
| GalN | - | - | - | 4 |

**Supplementary Table 8**. **Spectral deconvolution of 1D ^13^C CP spectra of R. delemar cell walls** **at 4 h of germination under different labeling conditions**. unLCLM represents unlabeled conidia grown in labeled medium; LCunLM represents labeled conidia grown in unlabeled medium; and LCLM represents labeled conidia grown in labeled medium. Deconvolution was performed from 110 to 79 ppm using DMfit software. The C1, C3, and C4 carbon sites were used to estimate the relative contributions of chitosan (Cs), β-1,3-glucan (B), and chitin (Ch), respectively.

| Polysaccharides | ppm | Line width | T2 | Amplitude | Integral | % |
| --- | --- | --- | --- | --- | --- | --- |
| unLCLM | | | | | | |
| Ch1/B1 | 104 | 1.07 | 1.48 | 3847400 | 1.25E+09 |  |
| B3 | 87.2 | 1.7 | 0.93 | 1567776 | 8.21E+08 | 13 |
| C4 | 83.54 | 2.58 | 0.61 | 1370045 | 8.15E+08 | 13 |
| Cs1 | 102.17 | 1.6 | 0.99 | 1161000 | 5.62E+08 | 74 |
|  | 101.28 | 2 | 0.79 | 1200500 | 7.20E+08 |  |
|  | 98.53 | 2.7 | 0.59 | 3507800 | 2.81E+09 |  |
| LCunLM | | | | | | |
| B1/Ch1 | 103.83 | 0.71 | 2.23 | 3695729.27 | 8.11E+08 |  |
| B3 | 87.01 | 0.92 | 1.72 | 1838329.11 | 5.27E+08 | 41 |
| Ch4 | 83.17 | 1.92 | 0.82 | 159660.66 | 9.35E+07 | 26 |
| Cs1 | 99.47 | 1.53 | 1.04 | 187331.02 | 8.76E+07 | 33 |
|  | 98.27 | 1.35 | 1.17 | 178107.57 | 7.41E+07 |  |
|  | 102.26 | 2.46 | 0.64 | 130510.67 | 9.50E+07 |  |
|  | 100.94 | 1.96 | 0.81 | 178107.57 | 1.06E+08 |  |
|  | 98.82 | 1.86 | 0.85 | 178107.57 | 1.01E+08 |  |
| LCLM | | | | | | |
| B1/Ch1 | 103.83 | 1.2 | 1.32 | 5561361.53 | 2.03E+09 |  |
| Ch4 | 83.52 | 2.45 | 0.65 | 1199440.42 | 8.16E+08 | 15 |
| B3 | 87.1 | 1.32 | 1.2 | 2697473.93 | 1.08E+09 | 65 |
| Cs1 | 98.37 | 2.45 | 0.65 | 1067260.99 | 7.81E+08 | 21 |
|  | 101.24 | 2.45 | 0.65 | 714782.52 | 5.19E+08 |  |
|  | 99.9 | 2.53 | 0.63 | 802902.14 | 6.03E+08 |  |

**Supplementary Table 9. Experimental parameters for proton-detected ssNMR experiments.** All experiments were conducted on an 800 MHz (18.8 T) spectrometer at a MAS frequency of 40 kHz. Selected experiments were additionally acquired on a 600 MHz spectrometer using a MAS frequency of 60 kHz.

| Expt. | B_0_  (T) | CP (µs) | | D1 | NS | td2 | td1 | aq2  (ms) | aq1  (ms) | decoupling | RFDR  (ms) | Samples |
| --- | --- | --- | --- | --- | --- | --- | --- | --- | --- | --- | --- | --- |
|  |  | t_cp1_ | t_cp2_ |  |  |  |  |  |  |  |  |  |
| CP | 18.8 | 600  (HC-CP) | - | 3 | 1024 | 2352 | - | 19.9 | - | - | - | Melanin Inhibited  CW Fraction &  Conidia |
| 2D hCH | 18.8 | 600  (HC-CP) | 600  (CH-CP) | 3 | 16 | 1204 | 512 | 19.9 | 6.4 | slpTPPM  (rf 10 kHz) | - |  |
| 2D hCH | 14.4 | 1000  (HC-CP) | 50  (CH-CP) | 2 | 32 or 64 | 1600 | 192 | 14.9 | 5.3 |  | - | Conidia &  5 h germination |
| 2D hChH  (RFDR) | 14.4 | 1000  (HC-CP) | 500  (CH-CP) | 3 | 512 | 1600 | 192 | 13.6 | 3.2 | WALTZ-16 | 0.5 |  |

**Supplementary Table 10. ^1^H and ^13^C chemical shifts of *R. delemar* polysaccharides.** For each carbon site, the ^13^C and ^1^H chemical shifts are shown in the top. The C8 position corresponds to the N-acetyl methyl group in chitin. (Bottom) Long-range intermolecular correlations identified using ^1^H and ^13^C using 2D ^1^H-detected chemical shifts (ppm) and indicate spatial proximity between β-1,3-glucan (B), chitin (Ch) and chitosan (Cs).

| Biomolecule | C/H1 | C/H2 | C/H3 | C/H4 | C/H5 | C/H6 | C/H8 | References |
| --- | --- | --- | --- | --- | --- | --- | --- | --- |
| β-1,3-glucan | 104.0  4.9 | 74.5  3.6 | 87.3  3.6 | 68.6  3.3 | 77.7  3.3 | 61.5  3.8 | --- | Kang et al 2018^1^  Cheng et al 2024^3^  Yarava et al 2025^9^ |
| Chitin | 104.2  4.9 | 55.3  3.7 | 73.6  3.8 | 83.6  3.7 | 75.2  3.8 | 61.5  3.8 | 23.4  2.2 |  |
| Chitosan | 99.2  4.8 | 55.3  3.7 | 68.6  3.7 | 80.6  4.4 | 71.0  4.1 | 61.3  4.0 | --- |  |

**Supplementary Table 11. Intermolecular interactions between glucans in *R. delamar*.** Long-range interactions detected by **^13^C-^1^H s**howing correlations between carbohydrate components, including glucan-chitin/chitosan interactions.

| conidia | Chemical shift  (^13^C-^1^H) ppm | 5 h culture | Chemical shift  (^13^C-^1^H) ppm |
| --- | --- | --- | --- |
| B3-ChMe | 104.2-7.1 | B1-ChH^N^ | 104.5-8.8 |
| B1-ChH^N^ | 103.6-7.1 | - | - |
| B5-ChH^N^ | 77.8-7.1 | Cs5-ChH^N^ | 72.9-8.7 |
| B4-ChH^N^ | 74.3-7.1 | Cs2-ChH^N^ | 55.1-8.7 |
| Cs3-ChMe | 66.9-1.4 | Cs3-ChMe | 69.5-1.2 |
| Cs5-ChH^N^ | 74.3-1.4 | Cs5-ChMe | 71.3-1.2 |
| Cs2-ChH^N^ | 56-7.1 | Ch4-H^N^ | 84.2-8.7 |
| Ch-ChH^N^ | 74.3-7.1 | Ch5-ChH^N^ | 75.1-8.7 |
| Ch1-H^N^ | 103.6-7.1 | Ch2-ChH^N^ | 54.7-8.7 |
| Ch5-ChH^N^ | 70.7-7.1 | Ch1-ChH^N^ | 104.5-7.4 |
| Ch2-ChN^H^ | 56-7.1 | Ch5-ChMe | 75.1-1.2 |
